# Transition from a meiotic to a somatic-like DNA damage response during the pachytene stage in mouse meiosis

**DOI:** 10.1101/328278

**Authors:** Andrea Enguita-Marruedo, Marta Martín-Ruiz, Eva García, Ana Gil-Fernández, M. Teresa Parra, Alberto Viera, Julio S. Rufas, Jesus Page

**Affiliations:** Departamento de Biología, Facultad de Ciencias, Universidad Autónoma de Madrid, Madrid, Spain

**Author notes:** Current address: Department of Developmental Biology, Erasmus Medical Center, Rotterdam, The Netherlands (Andrea Enguita-Marruedo). Correspondence to: Jesus Page.

**Keywords:** Meiosis, Homologous recombination, Non-homologous end joining, DMC1, RAD51, 53BP1, γH2AX, XRCC4

## Abstract

Homologous recombination (HR) is the principal mechanism of DNA repair acting during meiosis and is fundamental for the segregation of chromosomes and the increase of genetic diversity. Nevertheless, non-homologous end joining (NHEJ) mechanisms also act during meiosis, mainly in response to exogenously-induced DNA damage in late stages of first meiotic prophase. In order to better understand the relationship between these two repair pathways, we studied the response to DNA damage during male mouse meiosis after gamma radiation. We clearly discerned two types of responses immediately after treatment. From leptotene to early pachytene, exogenous damage triggered the massive presence of γH2AX throughout the nucleus, which was associated with DNA repair mediated by HR components (DMC1 and RAD51). This early pathway finished with the sequential removal of DMC1 and RAD51 and was no longer inducible at mid pachytene. However, from mid pachytene to diplotene, γH2AX appeared as large discrete foci. This late repair pattern was mediated first by NHEJ, involving Ku70/80 and XRCC4, which were constitutively present, and 53BP1, which appeared at sites of damage soon after irradiation. Nevertheless, 24 hours after irradiation, a HR pathway involving RAD51 but not DMC1 mostly replaced NHEJ. Additionally, we observed the occurrence of synaptonemal complex bridges between bivalents, most likely representing chromosome translocation events that may involve DMC1, RAD51 or 53BP1. Our results reinforce the idea that the early “meiotic” repair pathway that acts by default at the beginning of meiosis is replaced from mid pachytene onwards by a “somatic-like” repair pattern. This shift might be important to resolve DNA damage (either endogenous or exogenous) that could not be repaired by the early meiotic mechanisms, for instance those in the sex chromosomes, which lack a homologous chromosome to repair with. This transition represents another layer of functional changes that occur in meiotic cells during mid pachytene, in addition to epigenetic reprograming, reactivation of transcription, expression of a new gene profile and acquisition of competence to proceed to metaphase.

## Author summary

DNA repair is critical for both somatic and meiotic cells. During meiosis, hundreds of DNA double strand breaks (DSBs) are introduced endogenously. To repair this damage, meiotic cells use a specialized version of the homologous recombination (HR) pathway that uses specific meiotic recombinases, such as DMC1, to promote repair with the homologous chromosome instead of the sister chromatid. This process is important to ensure chromosome segregation during meiosis and, as a side consequence, increases the genetic diversity of offspring. Nevertheless, under specific circumstances, meiotic cells can use other DNA repair mechanisms such as non-homologous end joining (NHEJ), which is error-prone. We investigated the response of mouse spermatocytes to increased DNA damage caused by gamma radiation, which is commonly used in cancer therapy. We found that the excess of DSBs produced by irradiation is processed by the meiotic HR recombination pathway in spermatocytes at the early stages of first meiotic prophase. However, this response is not inducible from the mid pachytene stage onwards. From this point on, spermatocytes rely on a response that shares many features with that of somatic cells. In this response, the NHEJ pathway is first used to repair DNA damage but is subsequently replaced by a HR mechanism that does not use DMC1. Instead, it relies only on RAD51, which is known to function in both somatic and meiosis cells and, contrary to DMC1, has a preference for the sister chromatid. This switch from a meiotic to a somatic-like response is accompanied by a conspicuous change in the epigenetic response to DNA damage, reinforcing the idea that a functional transition occurs in meiotic cells during the mid pachytene stage.

## Introduction

DNA damage response is one of the most critical processes for cell survival and proliferation. Of the different forms of DNA damage, double-strand breaks (DSBs) are by far the most harmful. DSBs can arise spontaneously as a consequence of exposure to physical and chemical agents or following replication errors. In somatic cells, two main mechanisms operate to repair DSBs [1]. Non-homologous end joining (NHEJ) is the most common mechanism, working in all phases of the cell cycle [2]. Although this pathway is quite efficient, it is also error-prone as it does not discriminate whether the two rejoined ends were the correct ones. In contrast, homologous recombination (HR) uses an intact DNA molecule as a template for repair, ensuring high fidelity of repair. However, this mechanism only acts when a DNA copy, usually the sister chromatid, is available, which only happens during the S/G_2_ phases of the cell cycle.

During meiosis, homologous chromosomes undergo a series of complex processes, including pairing and synapsis, recombination and segregation. Meiotic recombination is in essence a HR DNA repair mechanism that ensures the proper segregation of chromosomes during the first meiotic division and increases genetic diversity [3,4]. Although the molecular mechanisms mediating HR in somatic and meiotic cells are similar, there are a number of differences, including the way DSBs are generated and some of the molecular components that work in their repair. In meiosis, HR begins after hundreds of DSBs are endogenously induced by Spo11 endonuclease during the leptotene stage of the first meiotic prophase [3,5,6]. The ATM kinase and the MRN protein complex (comprised of MRE11, RAD50 and NBS1) function as damage sensors by recognizing DSBs [7–9]. The MRN complex, together with other proteins (e.g. CtIP, BRCA1, BLM, EXO1, DNA2), then eliminates the covalent attachment of Spo11 and performs a 5’ to 3’ resection of DNA on either side of the break, which forms 3’-protruding ends of single-stranded DNA (ssDNA) [10]. The newly produced ssDNA is covered by RPA, which protects it from degradation [7]. Then, the ATR-ATRIP (Ataxia Telangiectasia and Rad3-related and ATR-Interacting Protein) complex binds directly to the RPA-coated ssDNA, thus localizing the kinase ATR to DSBs [8].

After DNA resection, the recombinase proteins RAD51 and DMC1 replace RPA and form nucleoprotein filaments, allowing the ssDNA to invade the DNA double helix of the homologous chromosome [4,10]. Template choice for DSB repair is another specific feature of meiotic HR: it is tightly regulated to favor inter-homologue recombination and crossing-over, which ensure coordinated chromosomal disjunction at the first meiotic anaphase. DMC1, which is only expressed during meiosis, drives repair to favor non-sister chromatid donors, while RAD51, which is essential for both somatic and meiotic recombination, acts to favor sister-chromatid donors [6,11–13]. DNA contacts between homologous chromosomes can ultimately resolve as reciprocal or non-reciprocal recombination events, which lead to crossovers or gene conversion events, respectively.

Although HR is the main DNA repair pathway acting during meiosis, NHEJ can also be used [14]. The action of NHEJ mechanisms is apparently simpler. Classical NHEJ relies on the recruitment of the Ku70/80 complex and other regulatory factors, such as 53BP1, to the site of breaks to prevent DNA resection. This is followed by the incorporation of DNA-PKcs and DNA ligase IV, which reseals the break with the help of accessory factors such as XRCC4 [7,10]. In recent years, in addition to the classical NHEJ, a variety of alternative NHEJ pathways, which use additional biochemical components, have been uncovered [10,15]. Components of the classical NHEJ pathway such as Ku70/80 and 53BP1 have been detected in meiotic cells, both in the course of normal meiosis [16] and after the exogenous induction of DNA damage [14,17]. Not surprisingly, this mechanism seems to be triggered only in the late stages of first meiotic prophase. This may be a consequence of the upregulation of HR repair during the early stages of meiosis following the endogenous production of DSBs by Spo11 and the resection of DNA that is concomitant with Spo11 removal [18]. However, coexistence of HR and NHEJ is possible during the late stages of meiotic prophase to repair DNA damage that was induced by either endogenous or exogenous mechanisms. Radiation exposure experiments, for instance, have reported an increase of both 53BP1 and RAD51 levels in pachytene and diplotene spermatocytes [14,17,19,20].

The key event for the choice between HR and NHEJ relies on the resection of DNA around the break [2]. The production of ssDNA overhangs hampers the action of NHEJ mechanisms, which require intact ends. Although the regulation of DNA resection at DSBs is not completely clear, a reciprocal regulation of factors promoting and inhibiting resection has been reported [21]. For instance, 53BP1 has been proposed to play a key role in inhibiting resection by hampering the loading of the CtIP-BRCA1 complex to the DNA, thus diverting repair to the NHEJ pathway [10,22,23]. CtIP-BRCA1, in turn, is thought to negatively regulate 53BP1 by inducing displacement of both 53BP1 and Ku70/80 from the break point and stimulating DNA resection by the MRN-(EXO1-DNA2-BLM) complex [2,24]. Interestingly, both 53BP1 and BRCA1 seem to rely on ATM kinase for phosphorylation, which is necessary for their function. Additional factors, such as the action of specific CDK-cyclin complexes and the epigenetic landscape around the break point also contribute to the regulation of DNA end resection [21].

In addition to the biochemical interactions described above, the morphological, temporal and epigenetic scenario in which DNA repair occurs during meiosis must be considered. Synapsis, the intimate association of homologues, is mediated by a highly specialized structure called the synaptonemal complex (SC) [25]. Assembly and disassembly of the SC during the first meiotic prophase is a tightly regulated process crucial for proper chromosome recombination and segregation [26], as evidenced by the number of synapsis mutants in which recombination is disturbed, and vice versa [27–31]. Furthermore, during first meiotic prophase, the complex regulation of transcription and chromatin modifications can influence the response to DNA damage [32–34]. Most conspicuously, histone H2AX is phosphorylated to give rise to γH2AX, which localizes throughout the nucleus during the leptotene stage in response to DSBs [35]. This contrast with the pattern of γH2AX in somatic cells, where it usually forms small and discrete foci [36]. γH2AX is involved in recruiting many DNA repair factors [5,10,35,37,38] and in the transcriptional silencing that characterizes the beginning of meiosis and sex chromosomes [33,35,39]. Notably, ATM, ATR and DNA-PKcs can all phosphorylate H2AX [10,40,41]; therefore, γH2AX is a marker of both the HR and NHEJ pathways. Upon DNA repair, γH2AX seems to be displaced from the chromatin and/or dephosphorylated by protein phosphatases [42–44].

To shed light on the complex relationships of DNA repair mechanisms acting during meiosis, we assessed DNA repair responses during mammalian male meiosis after the exogenous production of DSBs. We irradiated mice with gamma rays and then analyzed the localization and dynamics of various markers of DNA repair response, including γH2AX, DMC1, RAD51, 53BP1, Ku70 and XRCC4, at different times of recovery. We have uncovered two distinct epigenetic patterns in response to DNA damage in early and late prophase-I spermatocytes: a typical meiotic one and a somatic-like one acting at early and late stages, respectively. The transition to a somatic-like response during mid pachytene coincides with the sequential cessation of the meiotic HR response at mid pachytene and the consecutive activation of NHEJ and somatic HR repair mechanisms. In addition, we report the formation of chromosome bridges between non-homologous chromosomes associated with either HR or NHEJ markers.

## Results

### γH2AX dynamics after irradiation

We first analyzed the distribution pattern of γH2AX (H2AX phosphorylated at serine 139) in response to DNA damage. Phosphorylation of this histone is one of the first cytological events detected after DNA damage and has been used extensively to localize DSBs in both somatic and meiotic cells [5,10,35,36,38]. Staging of spermatocytes during first meiotic prophase was made on the basis of chromosome synapsis between autosomes and the morphology of the sex chromosomes following SYCP3 immunolabeling, as previously characterized [33].

In control spermatocytes, γH2AX is first detectable at early leptotene, when short threads of SYCP3 mark the initial assembly of axial elements (AEs) along the chromosomes. At this early stage, only a few discrete γH2AX foci are observed scattered throughout the nucleus (Fig 1A). During mid to late leptotene, when AEs form longer filaments, γH2AX is broadly localized throughout most of the nucleus (Fig 1B). This broad nuclear distribution is maintained during early zygotene (Fig 1C), when AEs start to synapse. From mid zygotene onwards, γH2AX signal decreases and, by the end of zygotene, is mainly associated with unsynapsed regions (Fig 1D). During pachytene, when homologous chromosomes are fully synapsed, γH2AX localizes almost exclusively on the sex chromosomes, which have extensive unsynapsed regions (Fig 1E and 2A-B). Nevertheless, large γH2AX foci are sometimes observed associated to the SCs of some autosomal bivalents. These foci have been previously described [5,33] and interpreted as unrepaired DSBs that tend to disappear with pachytene progression or, alternatively, as regions of transcriptional silencing [45]. During diplotene, when homologues desynapse, γH2AX remains present only on the sex chromosomes (Fig 2C).

**Figure 1.**
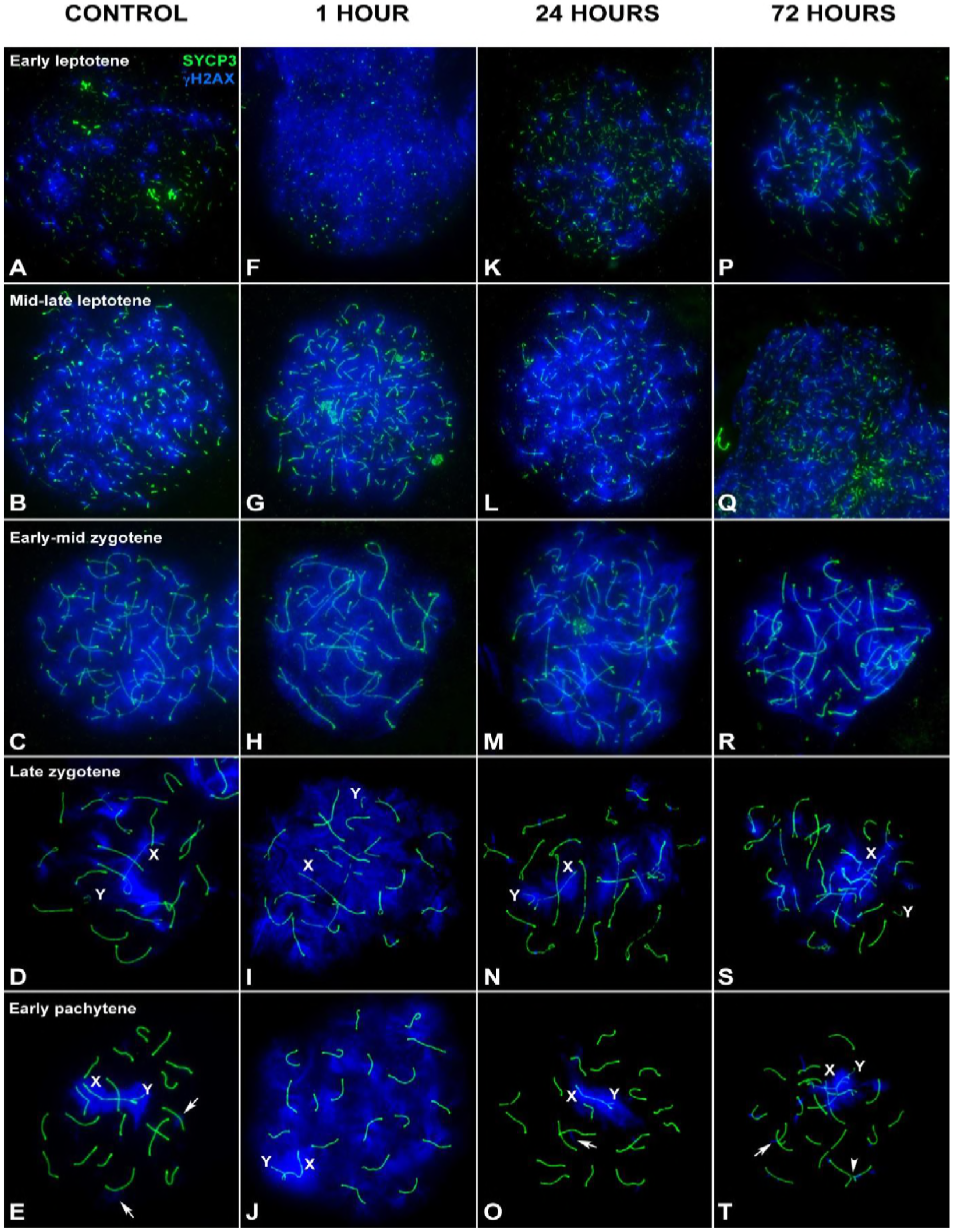
Pattern of γH2AX after irradiation in early prophase mouse spermatocytes. SYCP3 (green) and γH2AX (blue) at different stages of prophase I by recovery time after irradiation. (A-E) Control. (A) Early leptotene. γH2AX localizes as small scattered foci over the short threads of forming AEs. (B) Mid-late leptotene. AEs are more extended and now a massive γH2AX signal covers the nucleus. This pattern is also found in early zygotene (C), when AEs are completely formed and homologues start to synapse. (D) Late zygotene. Homologous chromosomes have nearly completed synapsis. γH2AX signal still occupies large chromatin regions, mostly on the unsynapsed autosomes and the X chromosome (X). The Y chromosome (Y) is usually devoid of massive γH2AX labeling. (E). Early pachytene. Autosomes are completely synapsed, whereas sex chromosomes show a variable degree of synapsis. γH2AX extends over both sex chromosomes (X and Y) and regions of chromatin around some autosomes (arrows). (F-J) 1 hour of recovery. Increased γH2AX signal is observed in the nucleus of spermatocytes from early leptotene to early pachytene. The signal covers the entire nucleus at all the stages, contrasting with the pattern of control cells. (K-O) 24 hours of recovery. There is an evident decrease in the amount of γH2AX in early leptotene, late zygotene and early pachytene spermatocytes, comparable to the controls. γH2AX localizes around the sex chromosomes, and some foci present in autosomes (arrows). (P-T) 72 hours of recovery. A pattern analogous to that at 24 hours is found. Chromosomal connections involving SYCP3 are observed between some bivalents (arrowhead).

**Figure 2.**
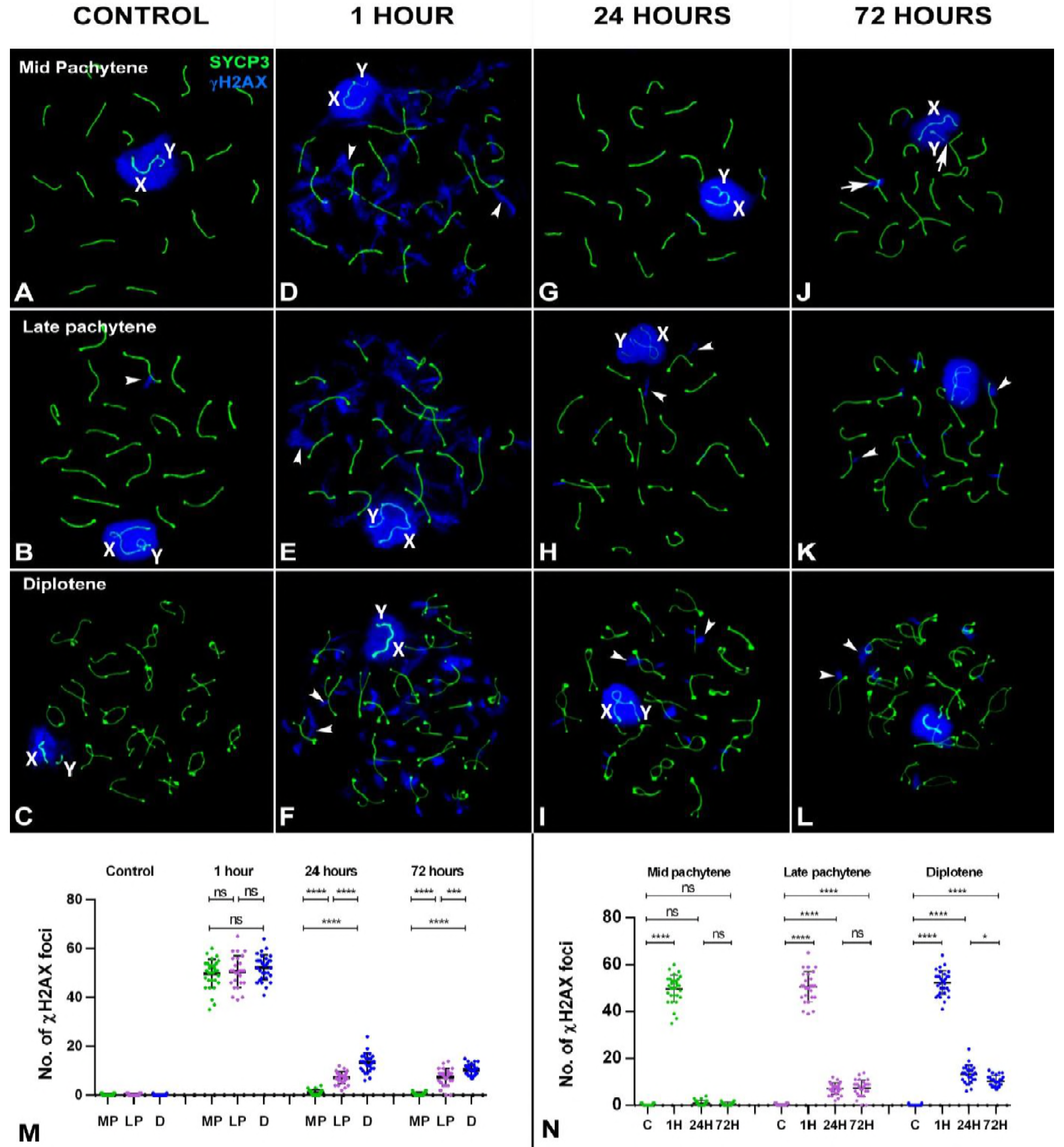
Pattern of γH2AX after irradiation in late prophase mouse spermatocytes. SYCP3 (green) and γH2AX (blue) at different stages of prophase-I by recovery time after irradiation. (A-C) Control. γH2AX appears around the sex chromosomes (X and Y) in mid pachytene (A), late pachytene (B) and diplotene (C). Occasionally, some small foci remain associated with autosomes (arrowheads). (D-F). 1 hour of recovery. In addition to the sex chromosomes, γH2AX localizes on the autosomes as large foci that emerge from the SCs (arrowheads) at all three stages. (G-I). 24 hours of recovery. All stages show a visible decrease in the amount of γH2AX. In mid pachytene cells (G), γH2AX foci are almost absent yet many foci are still present in late pachytene (H) and diplotene (I) cells. (J-L). 72 hours of recovery. The pattern is similar to 24 hours; some foci (arrowheads) remain present in late pachytene (K) and diplotene (L) cells. Some chromosomal connections are visible and appear to involve γH2AX signals (arrows). (M) Dotplot of the number of γH2AX foci in spermatocytes grouped by recovery time. The increase in the number of foci is evident 1 hour after irradiation. ANOVA analysis showed no statistical differences at this time between the three stages analyzed (p=0.22). Tukey’s multiple comparisons test for individual comparisons between different stages showed no statistical differences. 24 hours after irradiation, a reduction in the number of foci is observed at all stages, but now statistical differences between stages are observed (ANOVA p≤0.0001). Individual comparisons indicate the existence of differences between all stages. An analogous situation is found 72 hours after irradiation (ANOVA p≤0.0001). (N) Dotplot of the number of γH2AX foci in spermatocytes grouped by stage. While mid pachytene cells return to levels similar to the control 24 hours after irradiation (p≥0.05), late pachytene and diplotene do not at any time after irradiation. MP: mid pachytene; LP: late pachytene; D: diplotene; ns: non-significant; *: p≤0.05; **: p≤0.01; ***: p≤0.001; ****: p≤0.0001.

In gamma-irradiated spermatocytes, visible changes in the pattern of γH2AX localization are observed one hour after irradiation. In early leptotene cells, γH2AX is seen throughout the nucleus, in contrast to the small scattered foci seen in controls, indicative of a massive broadly distributed DNA repair response (Fig 1F). This pattern is also observed in late leptotene, zygotene and early pachytene spermatocytes (Fig 1G-J). Changes at late leptotene and zygotene stages are less evident as γH2AX is already broadly localized throughout the nucleus in control cells at these stages. This pattern indicates that cells at the beginning of meiosis up to early pachytene respond to the induction of DNA damage similarly. In contrast, the response of spermatocytes from mid pachytene onwards is rather focalized. Large γH2AX foci are observed emerging from the SCs (Fig 2D-F). These kind of signals have been called large foci [5], flares [45] or eruptions [46]. The morphology of these foci resembles that found in control spermatocytes (Fig 2B) and somatic cells [36]. These results reveal the existence of morphological differences in the response to DNA damage between early and late meiotic prophase spermatocytes.

Irradiated spermatocytes show a clear diminution of γH2AX in most stages 24 hours after treatment. Similar to control cells, early leptotene cells have a few scattered γH2AX foci (Fig 1K). If we consider that meiotic progression is not greatly affected by irradiation, then cells should progress to further stages during the recovery time. Therefore, these early leptotene cells could have been at preleptotene when irradiated (see S1 Figure for an estimation of the length of each meiotic stage, based on previous reports [47,48]). In late leptotene and early-mid zygotene spermatocytes, γH2AX is distributed throughout the nucleus, similar to control cells (Fig 1L-M). Likewise, the pattern of γH2AX at late zygotene and early pachytene is comparable to that of the controls (Fig 1N-O), in which γH2AX appears to label the unsynapsed chromosomal regions and some foci in a few chromosomes. These cells were likely irradiated at leptotene and zygotene stages, respectively, indicating that cells irradiated at early meiotic stages are able to achieve a control pattern corresponding to their stage 24 hours after irradiation. Contrastingly, cells from mid pachytene to diplotene retain several foci associated with SCs (Fig 2G-I). These differences are also observed 72 hours after irradiation (Fig 1P-T and 2J-L). In this case though, cells at mid pachytene 72 hours after irradiation were at an earlier pachytene stage at the time of irradiation. These cells likely had widespread localization of γH2AX at an earlier stage in response to DNA damage but their γH2AX pattern changes as they progress, very much like under endogenous production of DSBs. The persistence of γH2AX foci, however, indicates incomplete DNA repair.

Notably, leptotene cells were very scarce 72 hours after irradiation. Previous reports indicated that spermatogonia are particularly sensitive to radiation [17,19,49,50]. In order to confirm apoptosis of these cells, we performed a TUNEL assay on testicular sections (S2 Fig) and observed an increase of apoptosis in specific cell populations at different recovery times. Specifically, 24 hours post irradiation, a noticeable, but not massive, increase of apoptosis is observed in spermatogonia and prophase-I spermatocytes, while at 72 hours apoptosis is mainly observed in metaphase cells. This leads us to infer that irradiation may partially ablate spermatogonia population, but probably also interrupts the normal entrance of these cells in meiosis, which explains the scarcity of leptotene cells.

The two patterns of response to DSBs, early and late, also appear to differ in terms of γH2AX removal. Spermatocytes irradiated at late pachytene or diplotene, or those that reach these stages during recovery, remove γH2AX more slowly than those irradiated at earlier stages. In order to ascertain the efficiency of DNA repair, we recorded the number of γH2AX foci from mid pachytene to diplotene (S3 Table) and analyzed the progression of repair by recovery time (Fig 2M) and cell stage (Fig 2N). One hour after irradiation, the number of foci increases in the three stages. The ANOVA test showed no significant differences between stages at this time. However, at 24 hours, the number of foci returns to control levels in mid pachytene spermatocytes. In contrast, late pachytene and diplotene spermatocytes still show an increased number of foci, which is maintained even 72 hours after treatment. These results support the idea that γH2AX removal is less efficient as cells progress to later stages of prophase-I and that the number of foci seems to reach a steady state with no significant reduction.

### Formation of chromosomal bridges

One striking feature observed after irradiation is the formation of connections between non-homologous chromosomes, which can be visualized by SYCP3 immunostaining (Fig 1T, 2J and 3). Connections are only occasionally observed in control individuals (although in some mouse strains, they appear more frequently; unpublished results). Connections are observed at all post-treatment times (1, 24 and 72 hours) and could be clearly identified in zygotene to diplotene spermatocytes. On the basis of their morphological appearance, we classified connections in three categories (Fig 3): 1) distal contacts, in which chromosomes interact end-to-end (Fig 3A-B); 2) interstitial contacts, in which a filament emerges from one bivalent and contacts one or more bivalents laterally (Fig 3C-F) and 3) intrachromosomal contacts, in which the connection is observed within the same bivalent (Fig 3G-I). In some cases, the SYCP3-positive filament of a bivalent seems to split into two with a thin filament, probably involving a single chromatid, providing the connection (Fig 3C). In other cases, the filament appears thicker (Fig 3D).

**Figure 3.**
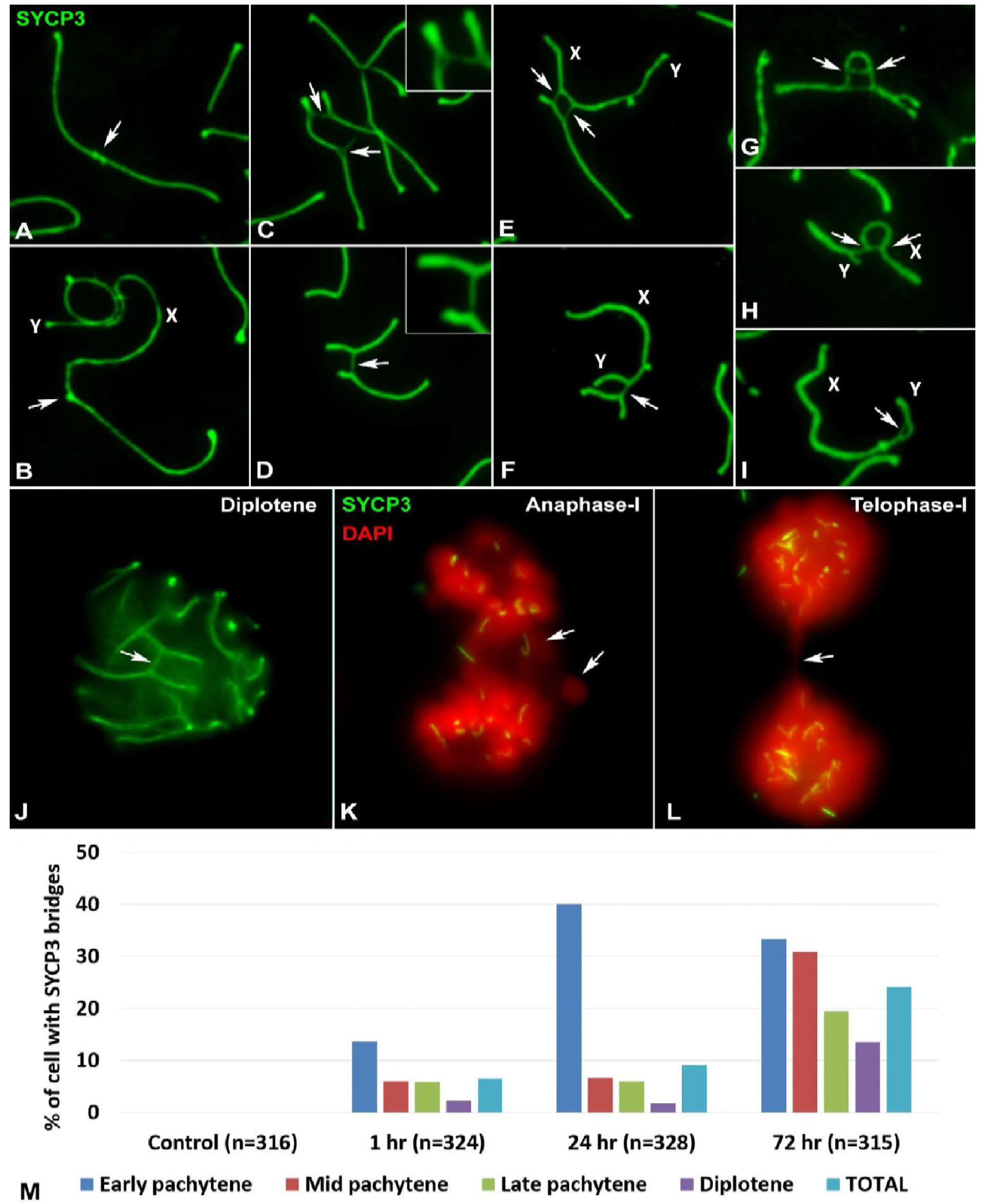
Types of chromosomal bridges. SYCP3 protein in green. (A-I) Spread spermatocytes at pachytene, except (G), which is at zygotene. (A) Distal junction between two autosomes. (B) Distal junction between an autosome and a sex chromosome, in this case the X. (C-D) Interstitial junctions between autosomal bivalents. In (C) a bivalent with two bridges, each contacting a different bivalent, is shown. In the inset, a higher power view of one of the bridges is shown. The lateral element of the homologue involved in the bridge is split into two filaments. One filament remains associated with the homologous chromosome and the other is linked to the chromosome of the other bivalent. In (D) two bivalents are sharing a bridge. In this case, the bridge is a whole counterpart, which has invaded the other bivalent. A higher power view of the bridge is shown in the inset. (E) Interstitial junctions between an autosomal bivalent and the X chromosome. In (F) a bridge is formed between an autosomal bivalent and the Y chromosome. (G) Chromosomal bridge within an autosomal bivalent. (H) Chromosomal bridge within the X chromosome. (I) Chromosomal bridge within the Y chromosome. (J-L) Squashed spermatocytes. DNA was counterstained with DAPI and false colored in red. (J) Bridges are seen in 3-dimension conserved cells. During anaphase-I (K) and telophase-I (L), chromosomal fragments and connections (arrows) are observed. (M). Graph showing the frequency of cells showing at least one bridge at the different cell stages of prophase-I and at the different time points after irradiation. The number of cells with bridges increases with recovery time. Chromosomal connections are especially represented in early pachytene cells. n = total number of cells analyzed.

Chromosome connections can be observed between autosomal bivalents, between autosomes and sex chromosomes or between sex chromosomes. The presence of these bridges is likely not an artifact of the spreading technique as they are also observed in squashed spermatocytes (Fig 3J). Furthermore, chromosome fragments and bridges are observed during anaphase- and telophase-I (Fig 3K-L), indicating that these connections may represent chromosomal translocations. While connections between bivalents can result in a non-homologous chromosomal translocation, bridges within bivalents can potentially link the two homologous chromosomes or different parts of the same chromosome. The presence of these chromosomal aberrations at metaphase-I, which are rarely detected in the control cells, might account for the increased apoptosis observed at this stage 24 and 72 hours after treatment (S2 Fig).

In order to understand the dynamics of chromosome bridge formation, we quantified the number of cells showing at least one of these chromosomal connections during pachytene and diplotene (Fig 3M) (connections were more difficult to discern from chromosome tangles in earlier stages). No bridges were found in the controls. However, after irradiation, the frequency of spermatocytes bearing bridges increases from 6.48% at one hour to 9.15% at 24 hours and 24.13% at 72 hours, indicating a clear rise in the number of bridges with time. Regarding the distribution of bridges by stage and time, at 24 hours, most of the cells with bridges are at early pachytene; however, by 72 hours, the distribution is more uniform among stages.

### Localization pattern of DMC1

In order to investigate the action of HR mechanisms, we first examined the spatial and temporal localization pattern of DMC1, which is exclusively present in meiosis and acts together with RAD51 [51,52]. To compare DMC1 distribution with the γH2AX pattern just described, we performed triple immunostaining of SYCP3, DMC1 and γH2AX. In control spermatocytes, DMC1 is first detected at the very beginning of leptotene, when AEs start to form along chromosomes. A few foci are seen scattered throughout the nucleus (Fig 4A), which are not specifically associated with either the short SYCP3 fragments or the small γH2AX foci already present. The presence of DMC1 foci at the beginning of leptotene suggested that they might be responding to DSBs produced by a SPO11-independent mechanism. However, their absence in SPO11 null mutants (S4 Fig) rules out this possibility. During late leptotene, many more DMC1 foci are clearly visible, and they appear to be mainly associated with chromosomal AEs (Fig 4B). During early zygotene, many DMC1 foci are still observed along both synapsed and unsynapsed chromosomal regions (Fig 4C). At late zygotene, the number of DMC1 foci decreases (Fig 4D). Some foci remain associated with autosomes but they are clearly more abundant on the unsynapsed AE of the X chromosome. A single DMC1 focus is observed on the Y chromosome. During early pachytene (Fig 4E), even fewer foci are visible. Although DMC1 and γH2AX are co-localized on some autosomes, in many cases, DMC1 and γH2AX foci are not associated with one another (see detail in Fig 4E), indicating that DMC1 localization persists after the removal of γH2AX. At mid pachytene, most autosomal DMC1 foci have disappeared, though the sex chromosomes still have a high number of foci (Fig 4F). DMC1 is no longer detectable at a cytological level after mid pachytene.

**Figure 4.**
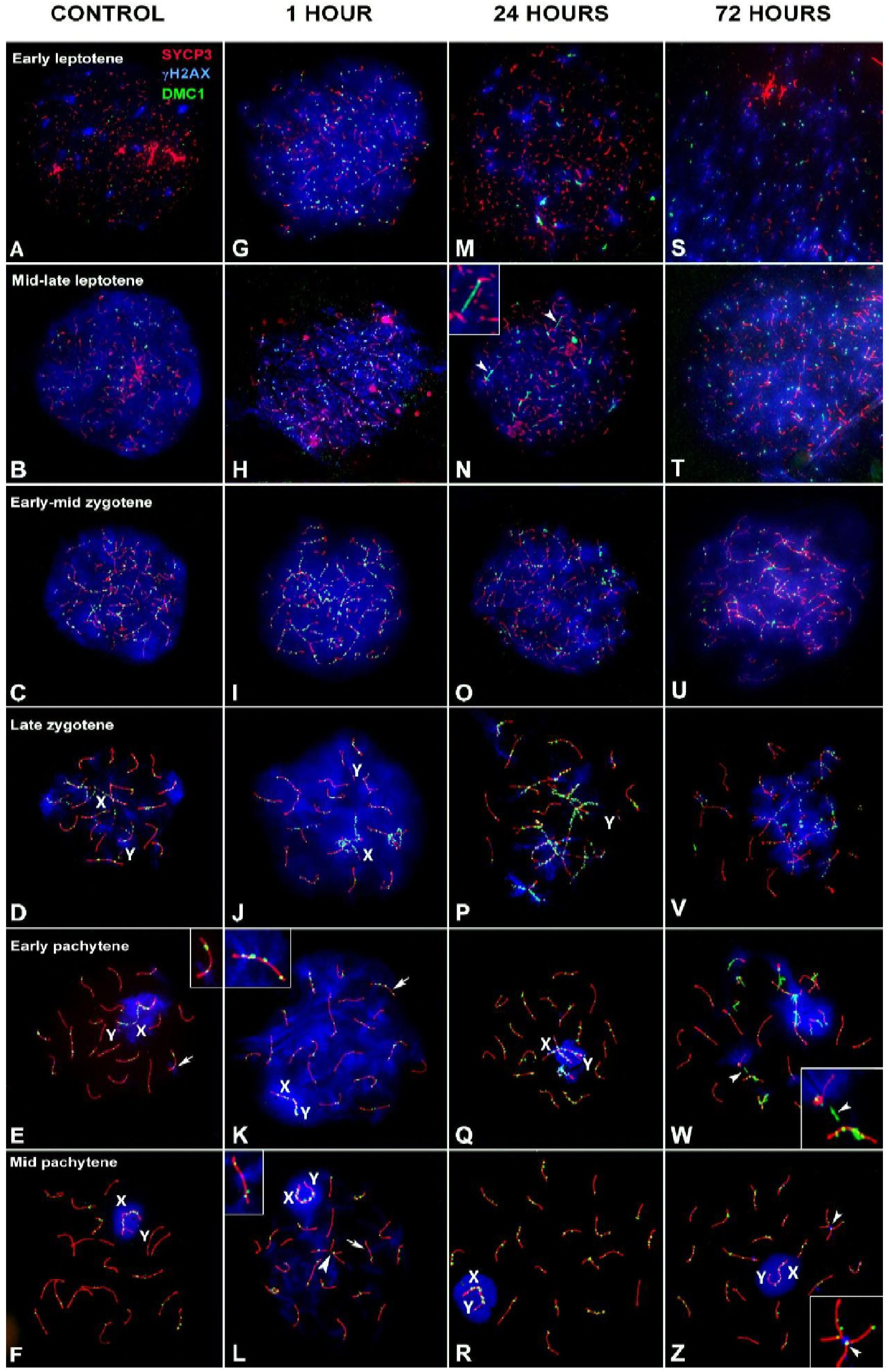
Pattern of DMC1 at different stages of prophase-I by recovery time after irradiation. SYCP3 (red), γH2AX (blue) and DMC1 (green). (A-F) Control. (A) Early leptotene. A few small foci of DMC1 appear distributed throughout the nucleus; however, they do not seem to specifically co-localize with SYCP3 or γH2AX. (B) Mid-late leptotene. DMC1 foci are more abundant and now mostly associated with short SYCP3 filaments. (C). Early-mid zygotene. DMC1 foci are very abundant over the formed AEs, some of which are undergoing synapsis. (D). Late zygotene. DMC1 foci are located over both synapsed and unsynapsed chromosomes. Some signal co-localizes with remaining clouds of γH2AX, while others do not. The X chromosome (X) appears coated with many foci, while only a single focus is seen on the Y chromosome (Y). Gradually, DMC1 foci disappear during early pachytene (E) and mid pachytene (F), but remain on the sex chromosomes and some autosomes. Only occasionally do some of these DMC1 foci co-localize with γH2AX (see detail in E). (G-L) 1 hour of recovery. The number of DMC1 foci increases at all stages from leptotene to early pachytene. At early leptotene (G), DMC1 coincides with the increase and spread of γH2AX to the whole nucleus. The number of DMC1 foci is clearly higher than in the control shown in A. No conspicuous differences in the pattern of DMC1 are observed at late leptotene (H) or zygotene (I-J). The Y chromosome still shows a single DMC1 focus. In early (K) and mid pachytene (L) spermatocytes, DMC1 is observed on autosomes and sex chromosomes. Again, DMC1 foci may co-localize or not with γH2AX. Enlarged views of some bivalents (arrows) are shown as insets in panels K and L. Chromosomal bridges are also found (arrowheads). (M-R) 24 hours of recovery. The distribution of γH2AX resembles that of control cells at all stages but the number of DMC1 foci seems reduced compared with cells 1 hour after irradiation. In some cells DMC1 appears to form filaments, sometimes joining AEs together (see arrowheads and detail in N). (S-Z). 72 hours of recovery. Leptotene cells (S-T) are found at a very low frequency and usually include morphological distortions. The morphological features of cells from zygotene to mid pachytene are similar to those found at 24 hours. Again, small DMC1 filaments appear on the chromosomes. These filaments seem to occasionally mediate the formation of bridges between two bivalents (arrowheads and details in W and Z); in some cases, γH2AX signal is associated with bridges (Z).

After irradiation, a notable increase in DMC1 protein expression is observed (Fig 4G-Z), with foci associating with unsynapsed AEs during leptotene, synapsed and unsynapsed regions during zygotene and synapsed autosomes and the AE of the X chromosome from pachytene onwards. DMC1 is not detected beyond mid pachytene, indicating that this protein is not inducible by radiation exposure after this stage. Similar to control cells, some co-localization of DMC1 and γH2AX is observed in irradiated pachytene cells (see details in Fig 4K and 4L). We also observed DMC1-positive filaments connecting different chromosomes. These filaments are mainly present at 24 and 72 hours after irradiation and likely represent the nucleoprotein filaments formed during the ssDNA invasion of the intact DNA copy. Although it is unclear whether these filaments join homologous or heterologous chromosomes at earlier stages (Fig 4N), by pachytene, heterologous associations are clearly observed. Indeed, some of these filaments appear to be associated with SYCP3 threads that bridge different bivalents (Fig 4W-Z), indicating a role for DMC1 in DNA repair between heterologous chromosomes under experimental conditions.

### Dynamics of DMC1 response

In order to analyze the dynamics of DNA repair associated with DMC1, we scored the number of foci in control and irradiated cells at different stages. On the basis of the morphological features of SC formation and the γH2AX localization pattern described above, we considered six different substages: early leptotene, mid-late leptotene, early-mid zygotene, late zygotene, early pachytene and mid pachytene. We did not record the number of DMC1 foci in leptotene cells 72 hours post irradiation given the scarcity of this cell population and the occurrence of morphological abnormalities, as mentioned above.

Our quantitative analysis revealed some interesting features (Fig 5 and S3 Table). First, the early leptotene cell population of control spermatocytes has a low number of DMC1 foci and very low standard deviation. As described above, this population is also characterized by a few small γH2AX foci. In contrast, mid-late leptotene cells show an increase in the number and standard deviation of DMC1 foci, in agreement with a previous report [53]. This stage is also associated with broad γH2AX labeling, as pointed above. Peak abundance of DMC1 foci occurs during early-mid zygotene and decreases thereafter. According to the ANOVA and Tukey’s multiple comparisons tests, differences between each stage and the next one are significant (Fig 5A), indicating that DMC1 distribution can be used to distinguish the cell populations of the six substages.

**Figure 5.**
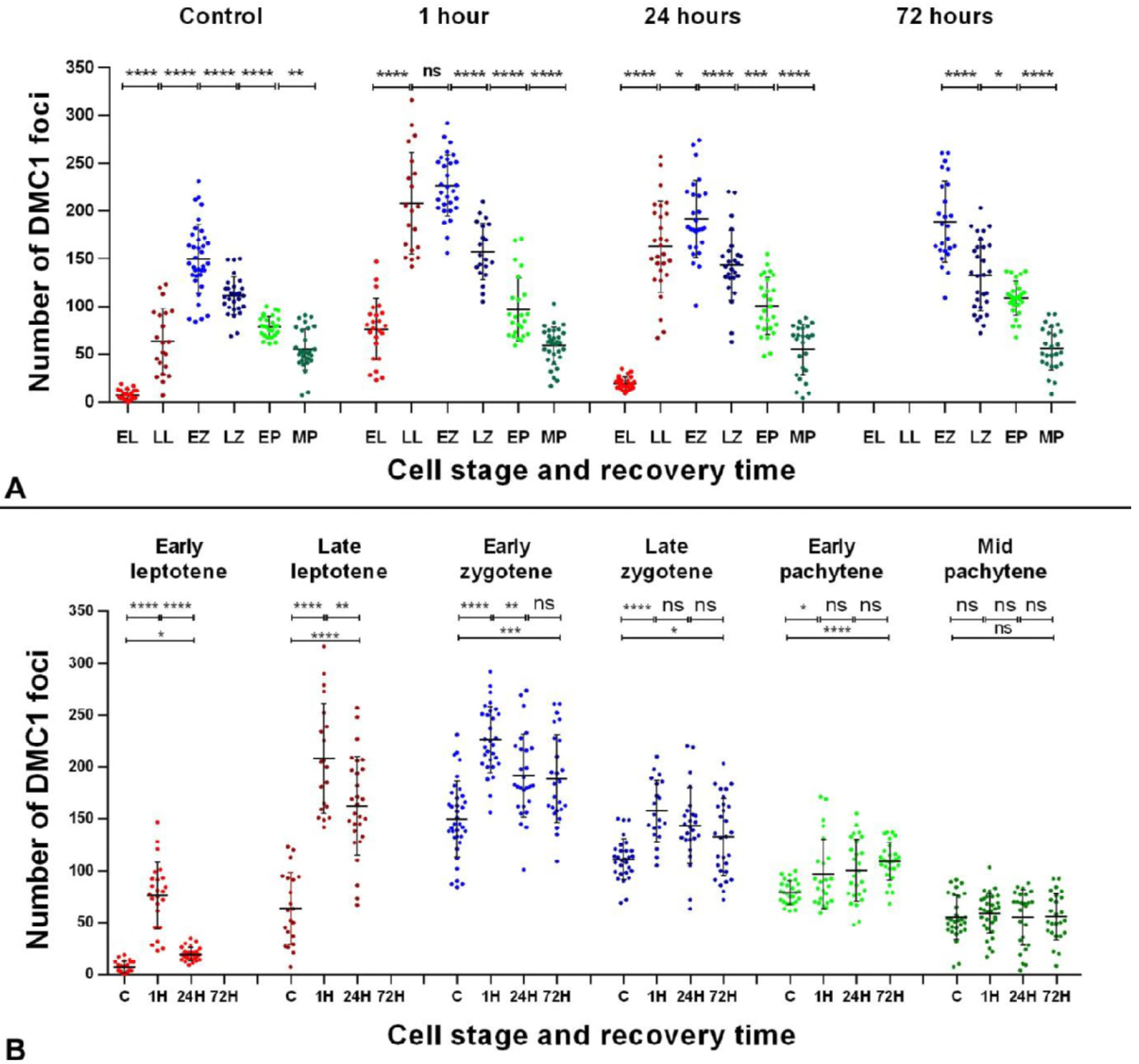
Dotplot representation of DMC1 foci distribution. (A) Analysis of DMC1 distribution by time of recovery. Six substages were considered (EL: early leptotene; LL: mid-late leptotene; EZ: early-mid zygotene; LZ: late zygotene; EP: early pachytene; MP: mid pachytene). The six populations, including early leptotene, are clearly distinguishable in the control. A low number of foci is found in EL cells but numbers increase in LL, peak in EZ and then gradually decrease in LZ, EP and MP cells. ANOVA analysis showed statistical differences (p≤0.0001) for the control and the three recovery times, and Tukey’s multiple comparisons test for individual comparisons between different stages showed statistical differences in all cases, except between LL and EZ 1 hour after irradiation (ns: non-significant; *: p≤0.05; **: p≤0.01; ***: p≤0.001; ****: p≤0.0001). (B) Analysis of DMC1 distribution by cell stage. ANOVA analysis showed that the number of DMC1 foci increased at all stages after irradiation (p≤0.0001) except mid pachytene (p=0.89). The increase of DMC1 foci observed 1 hour after irradiation compared to control is lower as cells are in more advances stages, and no increase is found at mid pachytene. Likewise, differences in the number of foci between 1 and 24 hours, or between 24 and 72 hours, become less or not significant as cells are in later stages. Nevertheless, control levels in terms of number of foci were not observed for any of the stages, even after 72 hours of recovery, except obviously mid pachytene. Tukey’s multiple comparisons test for individual comparisons (ns: non-significant; *: p≤0.05; **: p≤0.01; ***: p≤0.001; ****: p≤0.0001).

Second, as expected, the number of foci increases one hour after irradiation in most phases (Fig 5). As in the control, peak abundance of DMC1 foci is observed in early-mid zygotene spermatocytes, and each stage differs significantly from the following one, excepting mid-late leptotene and early-mid zygotene. However, the number of DMC1 foci induced by irradiation differs greatly among the different meiotic stages. The increase of foci compared to control is on average 69, 144, 76, 46, 18 and 4 for each of the six substages, respectively (see S3 Table). This striking result indicates that the cell stages are not equally sensitive to irradiation or that DMC1 localization to DSBs may be differentially regulated at the different stages due to the availability of this protein or other DNA repair factors. Furthermore, in irradiated mid pachytene spermatocytes, the number of DMC1 foci did not increase significantly regardless of recovery time, indicating that DMC1 is no longer inducible at this or later stages. These results can be more easily discerned when data are grouped by cell stage instead of recovery time (Fig 5B).

Third, after the increase of DMC1 foci immediately after irradiation, a slow diminution is observed with recovery time for most stages; however, most did not reach control levels even after 72 hours of recovery time (Fig 5B). Nevertheless, we observed two main stage-specific features. One, early leptotene cells show control levels 24 hours later. Moreover, while the number of DMC1 foci is quite variable one hour after treatment, 24 hours later, the range of foci narrows, very much like in the control. This finding may reflect the presence of newly formed leptotene cells that had just entered meiosis. Unfortunately, we could not record the number of DMC1 foci in early leptotene spermatocytes 72 hours after irradiation owing to the scarcity of this stage; and 2) in early pachytene, the number of DMC1 foci does not decrease but rather slightly increases with time. Assuming again that irradiation does not greatly disrupt meiotic progression, cells irradiated at a particular stage would continue to advance through meiosis and be at later stages when observed 24 or 72 hours later. Therefore, we arranged the quantitative data following a putative duration of 24 hours for leptotene, zygotene and early pachytene [34,47,50] (S1 and S5 Fig). This means that a cell irradiated at early leptotene would be at late leptotene-early zygotene 24 hours later and at early pachytene 72 hours later, and so on (S5 Fig). Considering four initial cell populations (early leptotene, late leptotene, early zygotene and late zygotene), we observed that, one hour after irradiation, the number of DMC1 foci increases in all cases and, in most cases, decreases 24 and 72 hours later, indicating efficient DNA repair in all cell populations. Nevertheless, control levels of DMC1 foci are not observed even after 72 hours of recovery, indicating that irradiation leads to an accumulation of DNA repair events. This contrasts with the quick and efficient removal of γH2AX at the same stages (Fig 1). The only exception are the cells that reach mid pachytene within the 72-hour period following irradiation. In this case, the levels of DMC1 did reach control levels, suggesting that DNA repair had been successfully completed in all cells. Alternatively, DMC1 might have been released from chromosomes at mid pachytene, regardless of whether repair had been completed or not.

### Localization pattern and dynamics of RAD51

We then analyzed the distribution of RAD51, which acts with DMC1 in the HR pathway, in control and irradiated spermatocytes. In agreement with previous reports [51,52], we found that RAD1 has a similar, albeit not identical, distribution pattern as DMC1 during first meiotic prophase (Fig 6). During zygotene stage (Fig 6A, S6 and S7 Fig), RAD51 localizes to the AEs of chromosomes, with a peak number of foci observed mainly in early-mid zygotene, decreasing continuously thereafter. At early pachytene and later stages, RAD51 foci remain associated with some autosomal SCs but are mainly found on the unsynapsed AE of the X chromosome (Fig 6B and 6C). Most of these foci are not associated with γH2AX, which at this stage is restricted to a few foci. RAD51 disappears during late pachytene (Fig 6D) and is absent at diplotene (Fig 6E). This pattern is very similar to that of DMC1; however, we observed that RAD51 remains associated with chromosomes for a longer period of time, into later pachytene stages. In order to observe this more clearly, we performed double immunostaining for both proteins (S6 and S7 Fig). During early stages of prophase-I, the localization of both proteins is almost, but not completely, identical. We noticed that not all DMC1 foci are associated with RAD51 foci and vice versa and that foci morphology can differ. More importantly, these two proteins are removed from chromosomes sequentially since the number of RAD51 foci on both autosomes and sex chromosomes clearly exceeds that of DMC1 at the mid to late pachytene transition (S6E-E” Fig). Thus, while the recruitment of RAD51 and DMC1 can be simultaneous upon the production of DSBs at the beginning of meiosis, persistent DSBs at the last stages of repair may lose DMC1 but maintain RAD51, which may reflect its role in promoting inter-sister versus inter-homolog interactions for the repair of DSBs [54].

**Figure 6.**
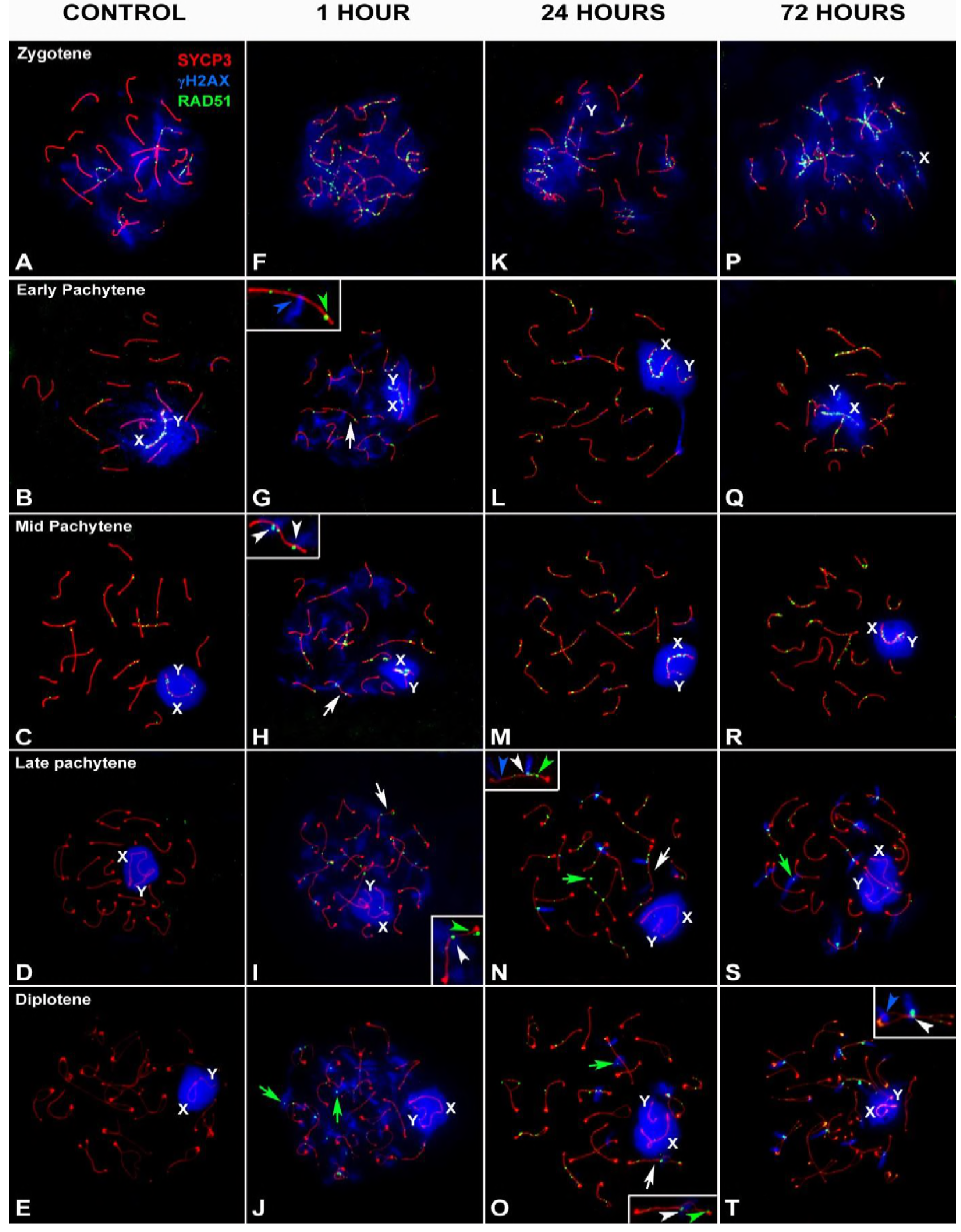
Pattern of RAD51 at different stages of prophase-I by different recovery time after irradiation. SYCP3 (red), γH2AX (blue) and RAD51 (green). (A-F) Control. (A) Late Zygotene. RAD51 foci are found in the non-synaptic AEs, which are also labeled with γH2AX, some synapsed autosomes and the sex chromosomes (X and Y). During early (B) and mid (C) pachytene, fewer RAD51 foci are observed. Some remain on the autosomes, but most are in the non-synapsed region of the X chromosome. RAD51 does not appear during the late pachytene (D) and diplotene (E). (F-J) 1 hour of recovery. Irradiation induces RAD51 in zygotene (F) and early pachytene (H) spermatocytes. RAD51 can still be detected in late pachytene (I) and diplotene (J) spermatocytes. Most RAD51 foci are located over the AEs or SCs (white arrows). Insets show enlarged views of RAD51 foci co-localizing with γH2AX (white arrowheads), RAD51 foci alone (green arrowheads) and γH2AXfoci alone (blue arrowheads). Some RAD51 foci are clearly detached from the AEs or SCs (green arrows in J). (K-O) 24 hours of recovery. RAD51 coincides with γH2AX during zygotene (K), while at later stages (L-O), co-localization of the two signals does not always occur (see detail in N). In late pachytene (N) and diplotene (O) spermatocytes, RAD51 foci are more abundant than at 1 hour. Foci are also larger. Green arrows indicate RAD51 foci not associated with SCs. (P-T) 72 hours of recovery. The pattern is similar to the results obtained after 24 hours of recovery. Most RAD51 foci are large and coincide with γH2AX during late pachytene (S) and diplotene (T).

Similar to the results with DMC1, we observed an increase in the number of RAD51 foci after irradiation (Fig 6F-T). Although we did not quantify the distribution of RAD51 at early meiotic stages, the broad co-localization of RAD51 and DMC1 up to mid pachytene (S6 and S7 Fig) suggests that both proteins follow a very similar pattern, i.e., peaking one hour after treatment then decreasing with recovery time. However, the localization patterns of these proteins are not identical. For instance, though RAD51-positive filaments bridging chromosomes are also observed (S7 Fig), they are thinner and scarcer than DMC1 ones.

Strikingly, after irradiation, RAD51 is observed in late pachytene and diplotene spermatocytes (Fig 6I,J,N,O,S,T). Given that RAD51 is not observed at these stages in control spermatocytes, these foci must represent newly localized protein induced after irradiation. Indeed, this RAD51 population differs with the one observed at earlier stages. First, the signal strength of RAD51 on the sex chromosomes is similar to that of autosomes. Second, foci tend to be larger and sometimes irregularly shaped. Finally, while virtually all RAD51 foci observed at early stages (up to mid pachytene) are associated with the AEs or SCs, during late prophase, a significant proportion of RAD51 is detached from the SCs (Fig 6J,N,O). Most of these foci co-localize with γH2AX one hour after irradiation, indicating they correspond to regions of DNA damage (Fig 6J and 6O).

To examine the dynamics of this late-appearing population of RAD51, we scored the number of foci present in mid pachytene to late diplotene spermatocytes (Fig 7A and 7B; S3 Table). This analysis uncovered some interesting features. First, one hour after irradiation, RAD51 increases significantly in all stages analyzed except mid pachytene. This striking result parallels the behavior of DMC1, which is also not inducible at mid pachytene, suggesting that HR repair can be compromised at this stage immediately after irradiation. The increase of RAD51 in late prophase spermatocytes is modest but significant, with no statistical differences observed among late pachytene and early and late diplotene stages. Second, after 24 hours of recovery, the number of RAD51 foci is significantly higher in all cell populations, though the increase is more pronounced in late pachytene cells and conspicuously lower in diplotene cells. After 72 hours of recovery, RAD51 levels remain high, though a slight decrease is observed in all stages, except mid pachytene. No statistical differences are detected between early and late diplotene, except in the number of foci at 24 hours. This difference could be caused by a small fraction of cells passing from late pachytene into early diplotene during this period, resulting in a higher number of RAD51 foci in early diplotene spermatocytes. The unexpected behavior of RAD51 during mid-late pachytene and diplotene stages suggests that the HR response to induction of exogenous DSBs initially may be absent or weak but increases with time, at least until 24 hours after irradiation.

**Figure 7.**
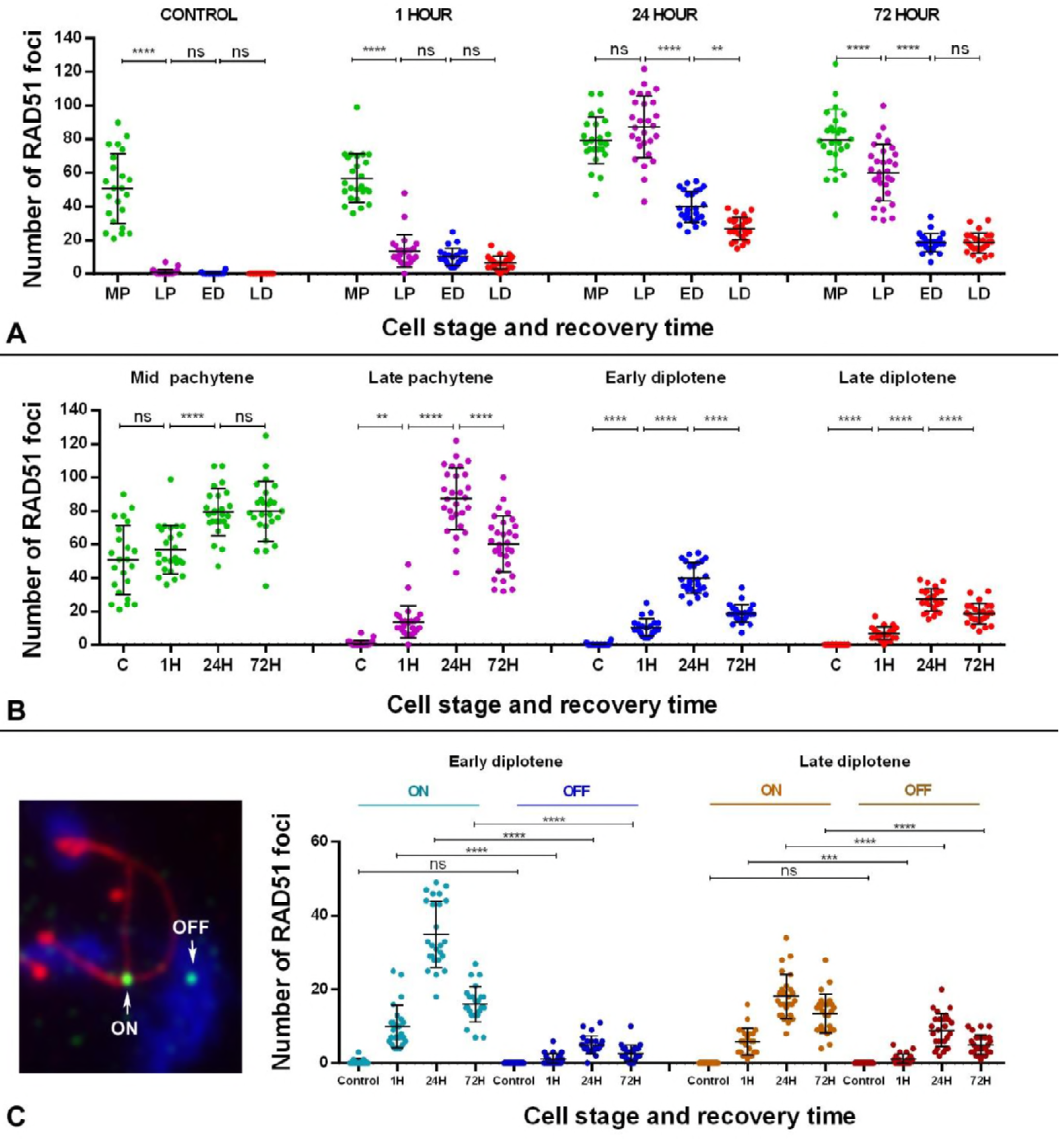
Dotplot representation of RAD51 foci distribution. (A) Analysis of RAD51 distribution by recovery time. Four substages were considered (MP: mid pachytene; LP: late pachytene; ED: early diplotene; LD: late diplotene). ANOVA analysis showed statistical differences (p≤0.0001) for the control and the three recovery times. In the control, MP cells have a high number of RAD51 foci but later-staged cells have little to none. A similar increase in the number of RAD51 foci is observed from LP to LD 1 hour after irradiation. Tukey’s multiple comparisons test for individual comparisons showed statistical differences between MP and the rest of the stages, and also between LP and LD. Twenty-four hour after irradiation, the increase in the number of RAD51 is more obvious at all stages, while 72 hours after irradiation, the number of foci decreases from LP to LD. (B) Analysis of RAD51 distribution by cell stage. The number of RAD51 foci increases significantly in cells at all stages after irradiation (p≤0.0001). However, according to Tukey’s test, the number of foci in irradiated mid pachytene cells 1 hour after irradiation is not significantly different from control cells. At 24 hours, the number of foci increases in mid pachytene cells and remains stable at 72 hours. This distribution departs from the pattern observed in cells at other stages, in which RAD51 increases slightly at 1 hour, peaks at 24 hours and then decreases at 72 hours. (ns: non-significant; *: p≤0.05; **: p≤0.01; ***: p≤0.001; ****: p≤0.0001). (C). Analysis of RAD51 foci associated (ON) or not associated (OFF) with SCs at early and late diplotene. The distribution of both kinds of foci is similar, increasing at 1 hour, peaking at 24 hours and decreasing at 72 hours. Notably, the proportion of foci not associated with SCs (OFF) is higher in late diplotene spermatocytes.

We were intrigued by the presence of RAD51 foci that were not associated with SCs. We analyzed the dynamics of RAD51 foci during diplotene (Fig 7C and S3 Table) and found that both SC-associated and non-associated RAD51 foci follow the same pattern, increasing one hour after irradiation, peaking 24 hours later and decreasing thereafter. We found that the number of foci associated with SCs is clearly higher in cells at early and late diplotene but that the proportion of non-associated RAD51 foci increases in late diplotene cells 24 and 72 hours after irradiation.

### Localization of NHEJ markers

In order to ascertain the action of the NHEJ repair mechanism, we studied the temporal and spatial localization of different components of this pathway. We first examined the localization of Ku70, which is involved in the protection of broken DNA ends, and XRCC4, which is a ligase-IV co-factor. Immunostaining of these proteins yielded nearly identical results; therefore, we will only show the localization of XRCC4 (Fig 8). Neither protein is observed during early meiotic stages in control spermatocytes (Fig 8A). At late pachytene, however, a weak signal appears throughout the nucleus (Fig 8B) and becomes more intense at diplotene (Fig 8C). At this stage, the signal appears slightly more intense over the sex chromosomes. In order to rule out the absence of XRCC4 labeling in early spermatocytes as an artifact of the spreading technique, we also immunostained testicular sections (S8 Fig). XRCC4 is absent in the basal spermatocytes of the seminiferous tubules and is only detectable in spermatocytes located in the middle of the epithelium, corresponding to late pachytene-diplotene cells. These results indicate that these components are present by default during the normal course of meiosis, in agreement with previous reports [14,16]. We observed a very similar pattern in irradiated spermatocytes: no signal is detected prior to late pachytene and, from this stage onwards, the proteins are distributed homogeneously throughout the nucleus (Fig 8D-L; S8 Fig). Although we did not quantify fluorescence intensity, no marked differences in the signal strengths of these proteins were observed between control and irradiated cells. Moreover, neither Ku70 nor XRCC4 accumulates at putative DSB sites after irradiation (e.g. in a pattern resembling that of γH2AX). Therefore, induction of DNA damage has little to no effect on the spatial and temporal localization of Ku70 and XRCC4, consistent with these proteins being present by default at these stages.

**Figure 8.**
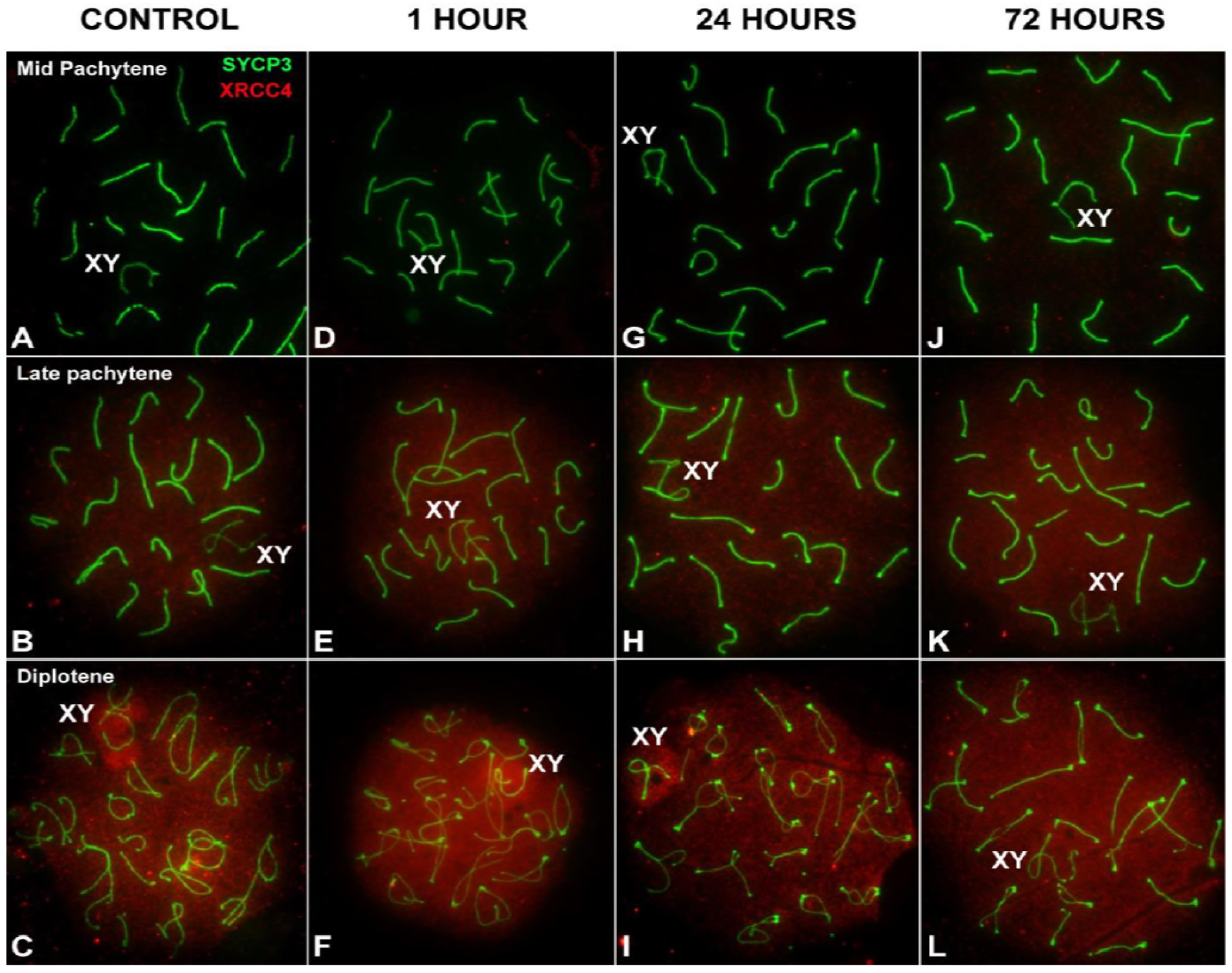
Distribution of NHEJ markers at different stages of prophase-I by recovery time after irradiation. SYCP3 (green) and XRCC4 (red) in late prophase-I spermatocytes. (A-C) Control. XRCC4 is absent up to mid pachytene (A). At late pachytene (B), a faint signal is observed in the nucleus, which becomes more intense at diplotene (C). The signal appears more concentrated on the sex chromosomes (XY). (D-F) 1 hour, (G-I) 24 hours and (J-L) 72 hours after irradiation. The localization pattern of XRCC4 at each stage is almost identical. Foci do not form at any stage or recovery time.

We also analyzed the localization of 53BP1, which has a main role in protecting broken DNA ends from resection during NHEJ repair. In control cells, 53BP1 is absent during leptotene, zygotene and early pachytene (not shown) but present by mid pachytene (Fig 9A), accumulating over the chromatin of the sex chromosomes in the same space occupied by γH2AX. 53BP1 signal is maintained during late pachytene (Fig 9B) and early diplotene (Fig 9C) but becomes weak by late diplotene. Occasionally, a weak signal is found on some autosomes.

**Figure 9.**
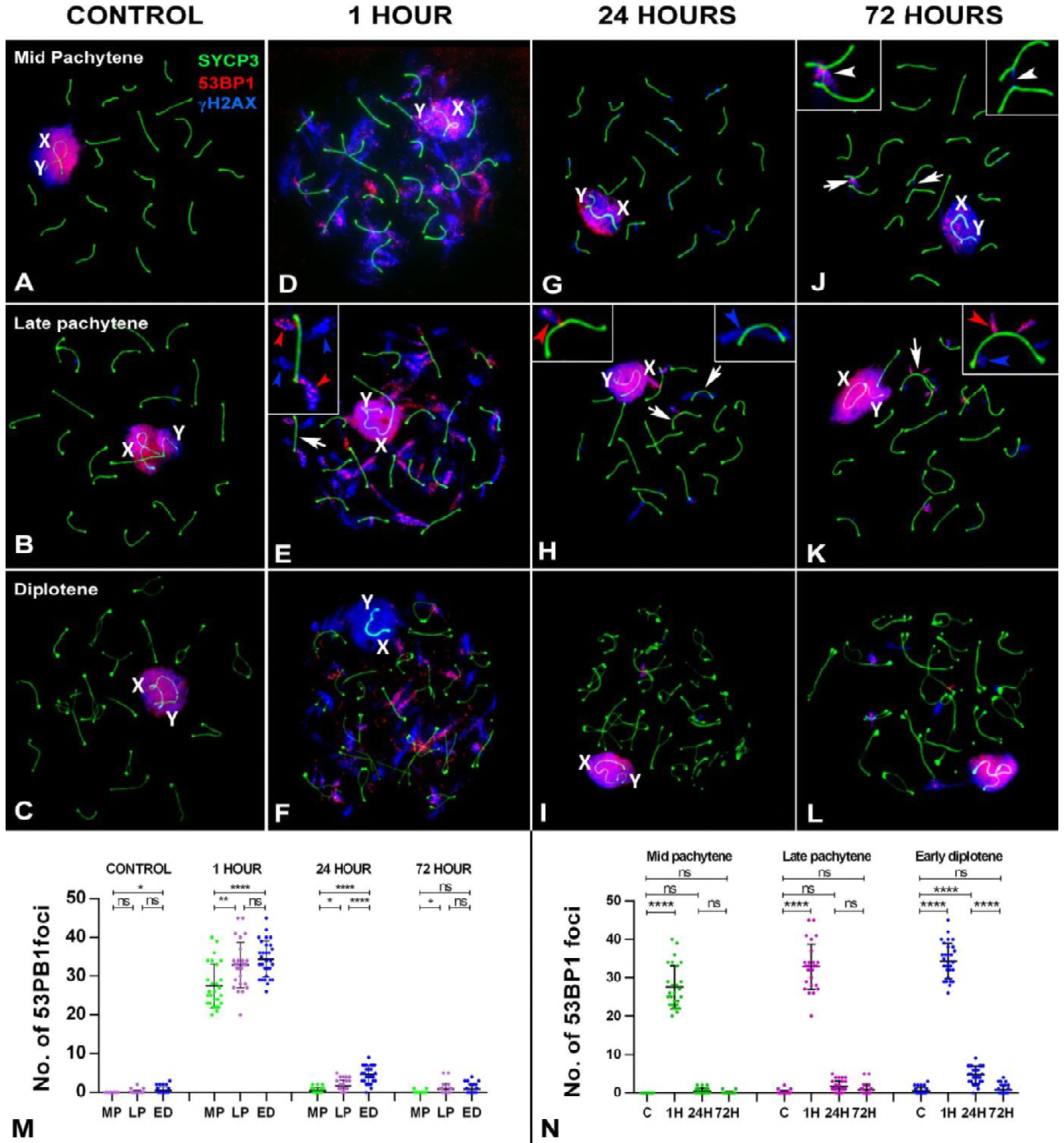
Pattern of 53BP1 at different stages of prophase-I by recovery time after irradiation. SYCP3 (red), γH2AX (blue) and 53BP1 (green). (A-C) Control. 53BP1 is first detected at mid pachytene (A) around the sex chromosomes and is maintained during late pachytene (B) and diplotene (C). During diplotene, the signal weakens, becoming no longer detectable by the end of this stage. The 53BP1 signal co-localizes with γH2AX around the sex chromosomes, X and Y. (D-F) 1 hour of recovery. From mid pachytene (D) onwards, a large number of 53BP1 foci appear on the autosomes as diffuse clouds. The 53BP signal is similarly maintained in late pachytene (E) and diplotene (F) spermatocytes, although foci become smaller as prophase-I progresses. Arrows indicate the bivalents shown in details. These 53BP1 signals largely coincide those of γH2AX (red arrowheads), although γH2AX foci without 53BP1 are also present (blue arrowheads) (see detail in E). (G-I) 24 hours of recovery. A noticeable decrease in the number of 53BP1 and γH2AX foci occurs relative to the 1-hour time point. In some cases, these foci coincide with those of γH2AX (red arrowhead in left detail in H) and in others they do not (blue arrowhead right detail in H). (J-L) 72 hours of recovery. The number and distribution of 53BP1 and γH2AX foci are similar to those at 24 hours. The presence of two interstitial bridges between autosomal bivalents (arrows) can be more clearly seen in the enlarged details in (J). γH2AX is observed on one of the bridges (right), whereas both γH2AX and 53BP1 are co-localized on the other (left). (M) Dotplot of the number of 53BP1 foci in spermatocytes grouped by recovery times. Three substages were considered (MP: mid pachytene; LP: late pachytene; ED: early diplotene). Increased numbers of foci are evident 1 hour after irradiation. ANOVA analysis showed statistical differences at this time between the three stages analyzed (p≤0.0001). Tukey’s multiple comparisons test for individual comparisons between different stages showed no statistical differences between LP and ED cells. A reduction is observed in the number of foci in cells at all stages 24 hours after irradiation. An analogous situation is found 72 hours after irradiation. (N) Dotplot of the number of 53BP1 foci in spermatocytes grouped by stage. Cells at all stages return to control levels 72 hours after irradiation. ns: non-significant; *: p≤0.05; **: p≤0.01; ***: p≤0.001; ****: p≤0.0001.

After irradiation, in addition to sex chromosomes, 53BP1 localizes to the autosomes from mid pachytene up to the end of diplotene. One hour after treatment (Fig 9D-F), a large number of irregularly shaped foci are observed on the autosomes, very similar to the γH2AX eruptions. Indeed, most 53BP1 foci on the autosomes co-localize with γH2AX, although unassociated foci of both proteins are also observed. The same pattern is found at both 24 (Fig. 9G-I) and 72 hours after treatment (Fig 9J-L). We observed that some chromosomal bridges, which are frequent in cells after treatment, are associated with 53BP1 (Fig 10J), indicating the involvement of NHEJ repair pathway proteins in this type of chromosome interaction.

**Figure 10.**
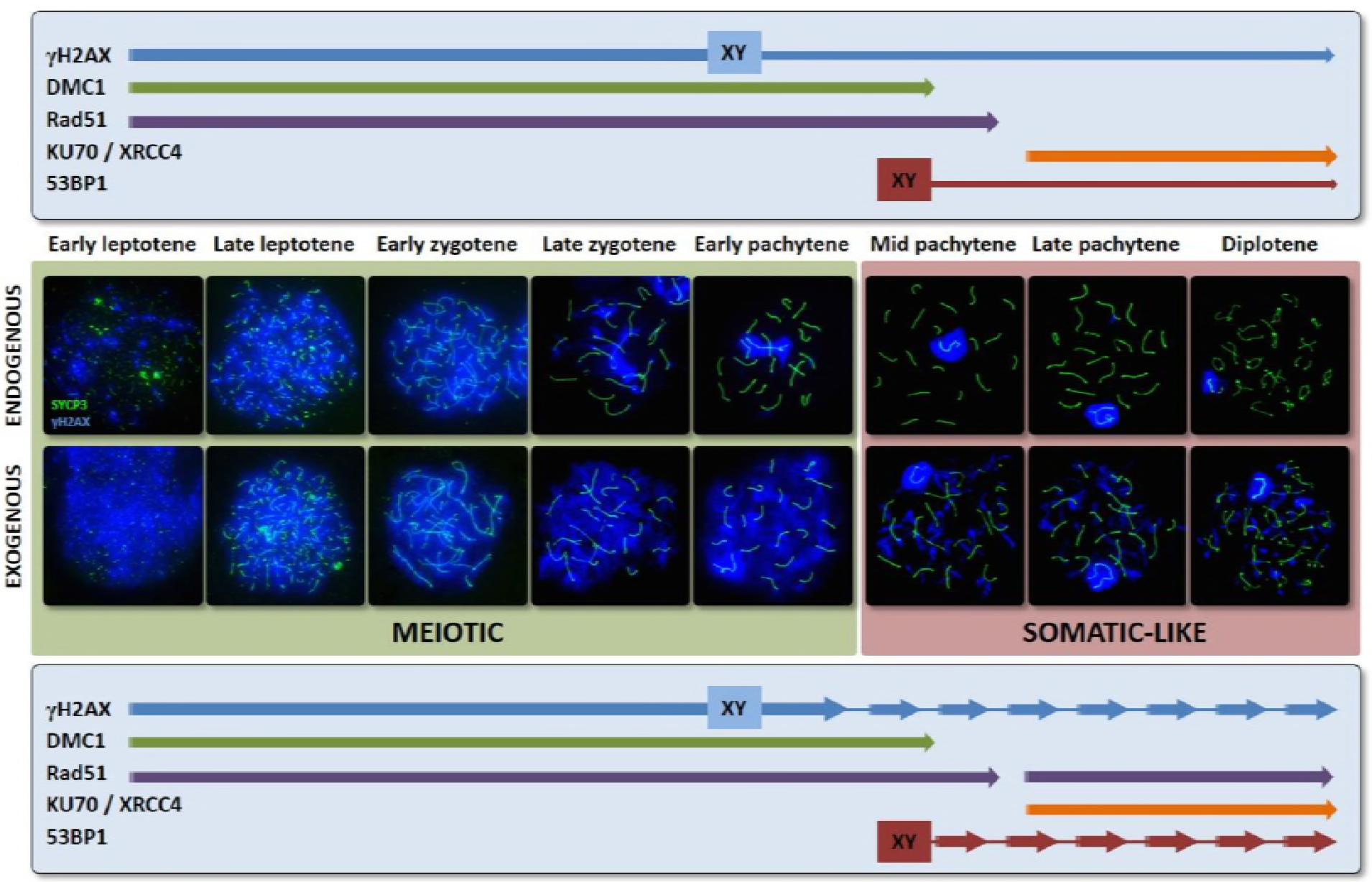
Model for the transition of the DNA damage response during meiosis. The early (meiotic) response works from early leptotene up to mid pachytene and is characterized by the action of HR mechanisms. This is the default pathway, likely due to the programmed resection of DNA upon SPO11 removal, which would hamper the action of NHEJ mechanisms. The meiotic response involves broad phosphorylation of γH2AX in the nucleus, likely in association with changes in chromatin organization, epigenetic modifications and transcriptional silencing, characteristic features of spermatocytes at these stages. DMC1 and RAD51 work together during this early response. DMC1 is removed first, leaving only RAD51 at the last stages of this response, which may affect interhomolog bias in the repair of DSBs. Induction of additional exogenous DSBs (but also potentially spontaneous, SPO11-independent ones) triggers an identical meiotic response, marked by the massive γH2AX localization throughout the nucleus and the increase of DMC1 and RAD51 in cells at all stages up to mid pachytene. Although γH2AX is quickly removed, many unresolved DNA damage intermediates accumulate even after long periods of recovery, indicating that this mechanism is not completely efficient. The dual late response very much resembles the response of somatic cells, including the appearance of discrete γH2AX foci. NHEJ is the first mechanism activated in this late somatic-like response, triggered soon after induction of DSBs from mid pachytene onwards. Some factors, like 53BP1, may already be present and localized on the sex chromosomes, with others (Ku70, XRCC4) appearing by default during late pachytene. This mechanism can quickly respond to DNA damage and, under normal conditions, likely resolve most, if not all, endogenously generated DSBs. However, after the induction of an exceeding number of DSBs, the initial NHEJ response is replaced by a HR one, involving only RAD51. Although this somatic-like response is less efficient in removing γH2AX than the early meiotic response, its overall repair efficiency is probably similar. Indeed, lower accumulation of unresolved intermediates is observed for this late response after long periods of recovery. The transition between these two DNA damage responses clearly occurs during mid pachytene, when the meiotic response is no longer inducible and the somatic-like one becomes available. This transistion indicates a possible physiological shift in meiotic cells as they prepare for further stages of first meiotic division and, more relevantly, chromosome segregation.

The quantitative analysis of 53BP1 after treatment (Fig 9M-N, S3 Table) shows the dramatic increase in the number of foci one hour after irradiation in mid- and late pachytene and early diplotene spermatocytes and its sharp decline 24 and 72 hours later. This pattern clearly contrasts with and seems antagonistic to that of RAD51, with NHEJ proteins acting as a fast response and HR proteins acting in two phases, weakly immediately after DNA damage and strongly 24 hours later. We also observed that late stages tend to have more 53BP1 foci, indicating stage-specific differences in the response. Nevertheless, by 72 hours after irradiation, all stages show control levels of 53BP1.

## Discussion

The accurate repair of DNA damage is critical for the survival of cells. Meiosis is an excellent model to investigate the response of cells to genomic damage owing to the occurrence of programmed DNA DSBs. However, the response to this endogenous damage must coexist with the sporadic occurrences of exogenous DNA damage, for instance, that caused by exposure to ionizing radiation. The specific processes that occur during meiosis, with the assembly of the SC being the most relevant, work in combination with the endogenous program to bias DSB repair towards the HR pathway [14,16]. Nevertheless, at the end of first meiotic prophase, some constraints might be relaxed, allowing the operation of somatic-like mechanisms. The results presented here offer new ways to understand the interplay of these two responses including how and when this transition occurs during meiosis.

### Different stages of prophase I have different responses to DNA damage

Phosphorylation of histone H2AX is one of the first key events to occur in response to DNA damage. As shown here and in previous reports, one hour after exposure to gamma radiation, γH2AX levels increase during all stages of first meiotic prophase [20]. However, two types of responses can be clearly distinguished according to the cellular phase: a massive response, characterizing the early stages, in which γH2AX marks the entire nucleus, and a more focused response from mid pachytene to diplotene in which γH2AX instead localizes as large and well-defined foci. This focused response is typically found in somatic cells [36] even though, under irradiation overexposure, both somatic and meiotic cells can show a pan-nuclear response [20,55]. However, in our case, all cells were exposed to the same dose of irradiation; therefore, the differences in response are not due to dosage-dependent effects. The origin of these differences could be related, in part, to changes in chromatin configuration and transcriptional activity, as previously suggested for somatic cells [56]. Highly dynamic replacement and modification of histones and proteins associated with chromatin are known to occur during prophase-I [33,34,57-59]. Mouse spermatocytes in early prophase-I are characterized by a widespread distribution of histone H3 monomethylated at lysine 4 and trimethylated at lysine 9, which are both related to chromatin compaction and transcriptional repression [33,34,60]. These modifications are lost or re-localized between early and mid pachytene, concomitant with other relevant epigenetic changes, such as the incorporation of histone H1t, which is related to the competency of cells to proceed to chromatin condensation stages [57], and a general reactivation of transcriptional activity, which is accompanied by the acetylation of histone H3 and other associated factors [33,34,61,62]. Therefore, the epigenetic changes occurring in meiotic cells at this stage likely act as regulatory factors modulating the DNA damage response.

Changes in chromosome organization may also play a role in the shift in the DNA damage response. In *C. elegans*, changes in both chromatin conformation and organization of the SC central element are proposed to be involved in the change in the DNA damage response during the mid to late pachytene transition [63,64]. Indeed, exogenous damage can lead to desynapsis of homologous chromosomes [64]. Although no dramatic remodeling of the SC occurs in mouse spermatocytes during this transition, the gradual shortening of the SC during pachytene, which results in longer chromatin loops, is a feature that potentially resembles such reorganization and thus may change the framework in which DNA repair proteins function.

An additional cause of this change may be related to the different kinases that promote H2AX phosphorylation. At least two rounds of H2AX phosphorylation dependent on two different kinases have been proposed to occur in mouse meiosis: the first during leptotene involving ATM and the second at the end of zygotene involving ATR [39,46]. Our efforts to corroborate this hypothesis by immunostaining for kinases, including ATM, ATR and DNA-PK, were unsuccessful. However, indirect proof can be inferred. In this sense, ATM kinase activity seems to produce an amplification loop in the phosphorylation of H2AX that extends up to several megabases beyond the DSBs [41], whereas the phosphorylation produced by ATR and DNA-PKcs entails a more focused response in which the signal is limited to areas close to DSBs [65]. Therefore, the two responses we observed with γH2AX may reflect a main role of ATM at the beginning of prophase-I and a higher activity of ATR and DNA-PKcs at later stages [46]. Interestingly, the response of somatic cells to irradiation, in which most DSBs are repaired by NHEJ [66,67], usually produces discrete foci of γH2AX in the nucleus [36,41], similar to those found in pachytene and diplotene spermatocytes. Therefore, it seems that early stages have a meiotic-specific γH2AX response, while late stages have a repair response more similar to somatic cells (Fig 10).

Based on γH2AX removal, the early response seems to be more efficient as cells irradiated at early stages return to control levels 24 after treatment. In contrast, cells irradiated at later meiotic stages retain a number of γH2AX foci for the duration of recovery. This contradicts findings that, on the basis of the removal dynamics of several repair proteins, suggested repair of DSBs induced at early stages of meiosis is slower than those occurring at later stages [19]. Therefore, it is important to be cautious with these interpretations. We found that many of the γH2AX foci observed at the different stages and recovery times are not associated with DMC1, RAD51 or 53BP1. The persistence of γH2AX in late stages may not be completely related to a delay in the completion of DNA repair but instead to delayed dephosphorylation or turnover of the histone [36]. A similar persistence of γH2AX foci has been also reported after etoposide-induced damage [20]. On the other hand, a substantial number of DMC1, RAD51 or 53BP1 foci are associated with chromosomes long after γH2AX has been displaced. Indeed, the number of DMC1/RAD51 foci in in cells irradiated at early stages is still above control levels 72 hours after irradiation (except in mid pachytene cells); this also applies for RAD51 and 53BP1 foci in late pachytene and diplotene cells, indicating the persistence of unrepaired events. Therefore, it seems that the production of exogenous DSBs challenges both early and late repair pathways during meiosis, resulting in an overall lower efficiency of meiotic repair of exogenous damage compared to somatic cells, as has been previously suggested [17,19].

### Response to endogenous DNA damage involves similar, but not identical, responses of DMC1 and RAD51

Our analysis of DSB repair pathways clearly indicates that HR is preeminent or exclusive at early meiotic stages, up to mid pachytene, and reveals interesting clues about the pattern of HR response under both normal and experimental situations.

In relation to the initiation of damage response in leptotene under normal conditions, we identified a population of early leptotene cells that is characterized by a low number of γH2AX and DMC1/RAD51 foci. Then, a burst of these proteins is detected in late leptotene. Although we cannot rule out the possibility that these two patterns are just the two extremes of a linear rise of DSBs during leptotene [53], it is also possible that they represent two different physiological stages. While early DSBs are clearly SPO11-dependent, the rate in which they arise is limited, probably owing to the action of a limited number of SPO11 complexes [53,68] or to restrictions imposed by associated factors. Some proteins that stimulate Spo11 activity, like IHO1, are associated with the AEs [69]. Therefore, DSB production could be limited in a chromosomal context in which AEs have not yet formed. Given this context, these DSBs would be able to only trigger a focus-limited (somatic-like?) γH2AX response. The extensive γH2AX labeling of this early leptotene population after irradiation indicates that these cells are competent to display a broad (meiotic) γH2AX reaction. Nonetheless, this *bona fide* meiotic response is only detected later in leptotene, once AEs have become more extended. This interpretation poses interesting questions about the transition from spermatogonia to meiosis, which includes other puzzling features like the premeiotic pairing of homologous chromosomes [70].

An intriguing issue arose when we compared the distribution of DMC1 and RAD51. We noticed that these two proteins tend to form mixed foci but that their localization patterns are not identical. Several studies reporting similar findings in budding yeast, plants and female mouse meiosis have suggested that these proteins occupy different positions along the nucleoprotein filaments [54,71,72], perhaps performing complementary functions. More strikingly, we found that the temporal pattern of DMC1 and RAD51 do not coincide, particularly in the transition from mid to late pachytene. DMC1 disappears from both autosomes and sex chromosomes at this stage, leaving only RAD51 on chromosomes. Temporal displacement between DMC1 and RAD51 loading and unloading has been also observed in plant meiosis [73]. This result may provide insight on the last stages of meiotic DNA repair pattern, particularly on the sex chromosomes. Several studies have hypothesized that DSBs on the X and Y chromosomes do not have homologous templates which can be used for repair, except obviously the pseudoautosomal region, and that repair can only be accomplished with the sister chromatid [32,35,74,75]. DMC1 may play a key role in interhomolog bias [54], such that its persistence on the sex chromosomes may explain why unresolved DSBs remain on these chromosomes long after most breaks have been repaired on the autosomes. The removal of DMC1 from sex chromosomes, and autosomes, at the mid-late pachytene stage may relax interhomolog bias, allowing RAD51 to then drive repair with the sister chromatid.

### Response to irradiation in early meiosis is characterized by the action of HR proteins that end at mid pachytene

Irradiation clearly stimulates the increase in the number of DMC1 and RAD51 foci from early leptotene up to early pachytene. We did not find any markers for NHEJ at these stages. Although we cannot rule out that alternative NHEJ pathways independent of Ku70 or 53BP1 might be present, it seems that the early response is mostly mediated by HR mechanisms, consistent with the findings of other studies [16,17,19]. It is reasonable to assume that induction of additional breaks will simply use machinery that is already present. Therefore, endogenous and exogenous DSBs can enter the same repair pathway. Consequences of this include, for instance, an increased number of chiasmata, as previously reported [19,76]. Nevertheless, this response is limited as prophase-I proceeds. The net increase of DMC1/RAD51 foci at each stage is lower as spermatocytes move to more advanced stages. Most strikingly, a meiotic HR response to exogenous DNA damage is very weak or not detected at mid pachytene, thus providing additional evidence of the functional shift of spermatocytes at this stage (Fig 10). This transition likely involves the cessation of expression of some meiotic-specific genes (like DMC1) and the initiation of a new gene expression profile [77–79]. This is interesting not only in terms of DNA repair but also in relation to the regulation of meiotic progression. Several studies have provided evidence of a pachytene checkpoint that monitors DNA repair, chromosome synapsis and other physiological processes such as sex chromosome inactivation [39,47,80–82]. Although the mechanisms that drive this checkpoint in mouse have not been completely elucidated, defective spermatocytes appear to be largely eliminated at a specific point of meiotic progression, identified as stage IV of the seminiferous epithelium in mouse, which most likely corresponds to the mid pachytene stage [47,80,83]. Once cells have cleared this checkpoint, inactivation of these surveillance mechanisms would be necessary to allow the progression of spermatocytes to later stages, allowing for instance desynapsis of chromosomes during diplotene without triggering meiotic progression arrest or inactivation of desynapsing regions. Likewise, inactivation of the early meiotic DNA damage response would be necessary after passing the checkpoint at mid pachytene, with new events of DNA damage that occur from this stage onwards being subject to new control mechanisms, as previously suggested [20]. These new mechanisms would rely on different checkpoints, as exemplified by the elimination of spermatocytes in mutant mice for late recombination proteins such as MLH1 or MLH3 at metaphase-I [84,85].

### NHEJ response is quickly stimulated upon irradiation from mid pachytene onwards

One may expect that, in the absence of a DMC1 response at mid pachytene, RAD51 takes the role of driving HR repair at this stage. As discussed above, RAD51 remains associated with chromosomes after DMC1 has detached, and previous reports have indicated that RAD51 is inducible in late prophase-I after irradiation treatments [17,19,20]. Consistent with this, one hour after irradiation, we found a modest but clear increase in RAD51 from late pachytene onwards; however, the increase was not significant at mid pachytene. Instead, we observed increased levels of 53BP1 at all stages from mid pachytene to late diplotene, indicating a faster response of the NHEJ pathway at these stages. Although RAD51 levels are increased in irradiated late pachytene and diplotene spermatocytes compared to controls, the number of 53BP1 foci clearly exceeds that of RAD51. This contrasts with a previous study that reported the presence of 53BP1 only after longer periods of recovery [17]. Differences in methodological approaches used to determine 53BP1 localization might account for these discrepancies.

The quick trigger of NHEJ in late prophase-I may be due to a change in the choice of the default mechanism for DSB repair (Fig 10). Somatic cells first attempt to use NHEJ to repair DSBs, even in the G2 phase of the cell cycle when a sister chromatid is available to carry out more reliable repair by HR [2,15,86]. The choice of NHEJ as the default mechanism from late pachytene onwards is illustrated by the constitutive presence of Ku70 and XRCC4 in the nucleus and the location of 53BP1 on the sex chromosomes. Therefore, as soon as new exogenous or even endogenous DNA breaks appear, this mechanism would quickly respond. This makes complete sense in terms of the biochemistry of DNA repair. Given that the choice between NHEJ or HR relies on the regulation of DNA resection around the break point [10], for which 53BP1 has an inhibitory role [22,23], it is clear that NHEJ must be a first option. Otherwise, once resection has been performed, repair by this mechanism would be no longer possible. Nevertheless, it also clear that both mechanisms are acting at the same time, raising the possibility that NHEJ and HR proteins are competing for DNA repair, especially from late pachytene onwards. In any case, after 24 hours of recovery, few 53BP1 loci remain, which may be due to rapid repair by NHEJ [2], but also to competition with HR. In somatic cells, HR seems to be preeminent in regions of high transcriptional activity [56], which is indeed the case of late pachytene and diplotene cells during meiosis. In any case, the increased presence of RAD51 foci 24 hours after irradiation indicates that HR repair mechanisms prevail again at that time.

### Late HR response involves RAD51 only

The late HR response has many differences with the early one. The most relevant is that it only involves RAD51. In the transition to a somatic-like DNA damage response, DMC1 is clearly no longer inducible, likely related to the change in gene expression pattern during pachytene [77–79]. The activity of RAD51 alone means that some of the constraints introduced by DMC1 in relation to DNA repair, such as homologous bias [6,11,12,54], would be relaxed. Therefore, this late repair could favor repair with the sister chromatid, which would be advantageous at the diplotene stage as desynapsis of chromosomes potentially hinders repair with the homologous chromosome.

The RAD51 foci present during the late HR response are larger. Although we do not have a clear explanation for the morphological change of RAD51 foci, differential organization of the repair machinery around the break point involving, for instance, the accumulation of several DSBs in each foci, or comprising the resection of longer DNA stretches, may account for this change, as previously suggested in *C. elegans* [63]. These foci are also correlated with the formation of smaller discrete γH2AX foci, which, as mentioned above, may be due to the action of ATR or DNA-PK over ATM in the phosphorylation of H2AX, leading to a different architecture of the repair foci [41,65].

Finally, the finding of RAD51 foci not associated with the AEs/SCs of chromosomes is an intriguing feature. Whether RAD51 foci are always associated with AEs/SCs has been a matter of long debate [87]. Current models propose that endogenous DSBs in early stages are produced either in the context of the AE or rapidly taken there by the action of regulatory factors, including MEI4, IHO1 and HORMAD1, among others [69,88]. This is probably provided by their ability to interact with SPO11 before or at the time of DSB production. We found that most DSBs induced after radiation also localize at the AEs/SCs, as revealed by the pattern of DMC1/RAD51 foci and as reported in a previous study [50]. Thus, at early meiotic stages, both endogenous and exogenously induced DSBs likely rely on similar mechanisms to be taken to the chromosomal axis. However, at late stages, the situation might be different. The presence of IHO1 and HORMAD1 has been reported in diplotene cells [69,89]; however, it remains unclear whether these proteins, or others required for DSB localization at the axes, are completely functional at these stages. Partial failure of this process might explain the fraction of DSBs located far from AEs/SCs. Likewise, the progressive loss of these proteins as prophase-I proceeds may account for the increased frequency of non associated foci in late diplotene spermatocytes compared with previous stages.

### Homologous recombination homeostasis and chromosomal bridges

The appearance of chromosomal bridges involving SYCP3-positive filaments is an intriguing feature that poses a number of questions about the nature of DNA repair. The presence of chromosomal connections and fragments is commonly found in irradiation experiments [90,91]; however, they are usually observed in metaphase and do not involve SC connections. To our knowledge, our study is the first to report that interactions between non-homologous chromosomes may involve not only the DNA contacts but also the axial structures they are attached to. This contrasts with the normal interactions of endogenous DSBs, which do not involve the formation of connections between the AEs of homologous chromosomes. A more in-depth characterization of these connections is needed to better understand the organization of the SC around break points and their role in promoting, facilitating or stabilizing chromosomal links.

Bridges appear soon after irradiation, indicating they are part of a very fast response, and at increased frequencies with recovery times. Moreover, although we cannot rule out that some bridges are present at leptotene (see Fig. 4N), they are undoubtedly present at all stages from zygotene to diplotene. In fact, according to our quantitative analysis, bridges are more frequent in early pachytene spermatocytes and less frequent in later phases. Several possible interpretations can be drawn from these results. Given that bridges are observed between non-homologous chromosomes, or between non-homologous sequences of the same chromosome (intrachromosomal junctions), one would expect that they correspond to DSBs repaired by NHEJ. However, the appearance of bridges at early stages such as zygotene and early pachytene, in which the main repair mechanism is HR, challenges this interpretation. Our immunostainings of Ku70, XRCC4 and 53BP1 indicate that NHEJ does not operate at early stages; however, alternative NHEJ mechanisms may potentially be present. Involvement of the NHEJ repair pathway in the formation of chromosomal bridges is doubtless only from mid pachytene onwards.

The presence of chromosomal bridges at early stages may be a consequence of the action of the HR pathway over homologous regions. This idea may seem completely counterintuitive but would be supported by the observation of DMC1/RAD51 filaments bridging non-homologous chromosomes. For HR to efficiently start, a minimum length of perfect homology is needed, which in mammals is 200-250 base pairs [10]. For this reason, repair templates are usually the sister chromatid or the homologous chromosome. However, the multitude of repeated sequences in the genome could provide sufficient homology to induce repair by HR. In the normal course of meiosis, endogenous DSBs are prevented over repeated DNA sequences. Moreover, mismatch repair mechanisms are in place to avoid recombination between highly homologous sequences of non-homologous chromosomes [92]. In addition, the number of DSBs is tightly regulated to not exceed a certain number, thus allowing mismatch repair mechanisms to function effectively [93]. Proteins like MEI4 and IHO1 seem to be involved in this limitation [69,88]. However, the excess of DSBs produced by radiation may have a deregulatory effect on the control of repair mechanisms such that homology requirements may be bypassed, thus allowing repair between non-homologous chromosomes.

Regardless of the mechanisms used to form bridges, the final output is the production of chromosome connections that likely lead to the occurrence of translocations and fragmentation. This may cause cells to be compromised in the faithful distribution of chromosomes during the first meiotic division. Indeed, a noticeable increase of apoptosis is observed in metaphase/anaphase cells 24 and 72 hours after irradiation.

## Conclusions

The results presented here provide new insights on the transition between different programs of DNA repair during meiosis that act in a stage-dependent manner. A switch in DNA damage repair responses during meiosis has been also reported in other animal models such as *C. elegans* [63,64] and *Drosophila* [94]. Strikingly, these transitions also occur at the mid or late pachytene stages, indicating they may represent a conserved feature of meiosis. However, since both *Drosophila* and *C. elegans* control synapsis and DNA repair differently than mammals, particularly as they lack a DMC1 orthologue, the regulation of this transition might be different.

The evidence presented also offers new clues about the location and dynamics of DNA repair mechanisms during meiosis and raises new questions about the differential functions performed by DMC1 and RAD51. The late HR pathway very much resembles the somatic response, presenting focalized γH2AX and involving only RAD51. This somatic-like response likely acts to repair DSBs that were not properly repaired by the meiotic default pathway (e.g. those on the sex chromosomes) or the occasional DNA damage that occurs after the primary phase of meiotic repair has concluded until cells start to condense chromatin and prepare for cell division. At this point, re-triggering a complex repair mechanism leading to the production of crossovers (provided chiasmata formation has been properly accomplished) would not be necessary. A simpler response using NHEJ or somatic HR would be sufficient. As previously suggested [63], this shift could simply be contributing to the maintenance of genome integrity before spermatocytes are engaged in segregating chromosomes to daughter cells.

## Materials and methods

Adult CD1 male mice were used in this study. Animals were kept at the animal facility of the Universidad Autonoma de Madrid, following the animal care standards of the institution. All experiments were approved by the UAM Ethics Committee (certificate CEI 55-999-A045). Males were exposed to 5Gy gamma radiation in a CIS Bio International irradiator, equipped with a Cesium^137^ source. Mice were sacrificed by cervical dislocation 1, 24 and 72 hours after irradiation and the seminiferous tubules processed as described below. Testicular samples of SPO11 knockout mice [31] were kindly shared by Dan Camerini-Otero (NIDDK, NIH, Bethesda, MD).

### Cell spreads and squashes

For spermatocyte spreads, we used the procedure described by Peters and coworkers [95]. Seminiferous tubules were disaggregated with forceps in a petri dish and a cell suspension was collected in phosphate buffered saline (PBS: 137 mM NaCl, 2.7 mM KCl, 10.1 mM Na2HPO4, 1.7 mM KH2PO4, pH 7.4). After tubule fragments settled to the bottom of the dish, the cell suspension was transferred to a tube and centrifuged. The pellet was then resuspended in 400 μl of 100 mM sucrose. Cells were spread onto a slide submerged in 1% formaldehyde in distilled water containing 50 mM Na2B4O7 and 0.15% Triton X-100 and then left to dry for two hours. Slides were subsequently washed with 0.04% Photo-Flo (Kodak) in distilled water and air-dried before being used for immunofluorescence or stored at −80°C.

Spermatocyte squashes were prepared as previously described [96]. Seminiferous tubules were fixed for 10 minutes in 2% formaldehyde in PBS containing 0.1 % Triton X-100. Fragments of tubules were placed on a slide coated with 1 mg/ml poly-L-lysine (Sigma) with two drops of fixative. A coverslip was put on top of the tubules and the cells were released by gently pressing the coverslip with a pencil. Finally, tubules were squashed, the slide was frozen in liquid nitrogen and the coverslip removed with a blade. Slides were immediately placed in PBS for further use.

### Immunofluorescence

Spread and squashed slides were rinsed three times for 5 min each in PBS and incubated overnight at room temperature with primary antibodies diluted in PBS. The following primary antibodies and dilutions were used: mouse monoclonal anti-SYCP3 (Abcam, 97672) at 1:200; rabbit anti-SYCP3 (Abcam, 15093) at 1:100; mouse monoclonal against histone H2AX phosphorylated at serine 139 (γ-H2AX) (Upstate, 05-636) at 1:1000; rabbit anti-DMC1 (Santa Cruz, SC-22768) at 1:50; rabbit anti-RAD51 (Santa Cruz SC-8349) at 1:50; rabbit anti-53BP1 (Abcam 36823) at 1:100; goat anti-XRCC4 (Santa Cruz, SC-8285) at 1:100; goat anti-Ku70 (Santa Cruz, SC-1486) at 1:50. After incubation, slides were rinsed in PBS three times for 5 minutes each and subsequently incubated with the appropriate secondary antibodies in a moist chamber at room temperature for 1 h. We used anti-rabbit, anti-mouse and anti-goat secondary antibodies raised in donkey and conjugated with either Alexa 350, Alexa 488, Alexa 594 (Invitrogen), DyLight 549 or DyLight 649 (Jackson ImmunoResearch). Slides were subsequently rinsed in PBS three times for 5 min each and mounted with Vectashield (Vector). For double detection of two antibodies raised in the same species, we used Fab secondary antibodies as previously described [97].

Observations were made on an Olympus BX61 microscope equipped with a motorized Z axis. Images were captured with an Olympus DP72 digital camera using the Cell-F software (Olympus, Hamburg, Germany) and processed using the public domain software ImageJ (National Institutes of Health, USA; http://rsb.info.nih.gov/ij) and Adobe Photoshop 7.0.

### Testicular sections and TUNEL assay

Testicles were fixed in cold 1% formaldehyde in PBS for 6 hours and then dehydrated and embedded in paraffin. Transverse sections (7 μm) were cut and mounted onto slides. Slides were then deparaffinized and treated with 0.1% sodium citrate buffer containing 0.1% Triton-X 100 for 10 min at 37°C. Sections were subsequently processed for immunofluorescence as described above or for TUNEL (Roche) following manufacturer instructions. Slides were counterstained with DAPI and mounted with Vectashield.

### Statistical analysis

γ-H2AX, DMC1, RAD51 and 53BP1 foci and chromosomal bridges were scored manually and the results were compared between different cell stages and times after irradiation. At least 25 cells were scored for each protein, treatment and cell stage. For the TUNEL assay, 300 tubules were analyzed for each treatment, recording the proportion of tubules with apoptotic cells and the total number of positive cells, which were classified as spermatogonia, prophase-I and division spermatocytes owing to their size, position on the seminiferous epithelium and chromosome condensation. Data were analyzed using ANOVA and Tukey’s multiple comparison tests for individual comparisons between different stages. Statistics and graphics were made using GraphPad Prism 6 or Excel software.

## Acknowledgements

We are grateful to Dr. Dan Camerini-Otero for generously sharing samples of SPO11 knockout mice and to Dr. Monica Pradillo and Willy Baarends for their critical review of the manuscript.

## Supplementary information

**S1 Figure.**
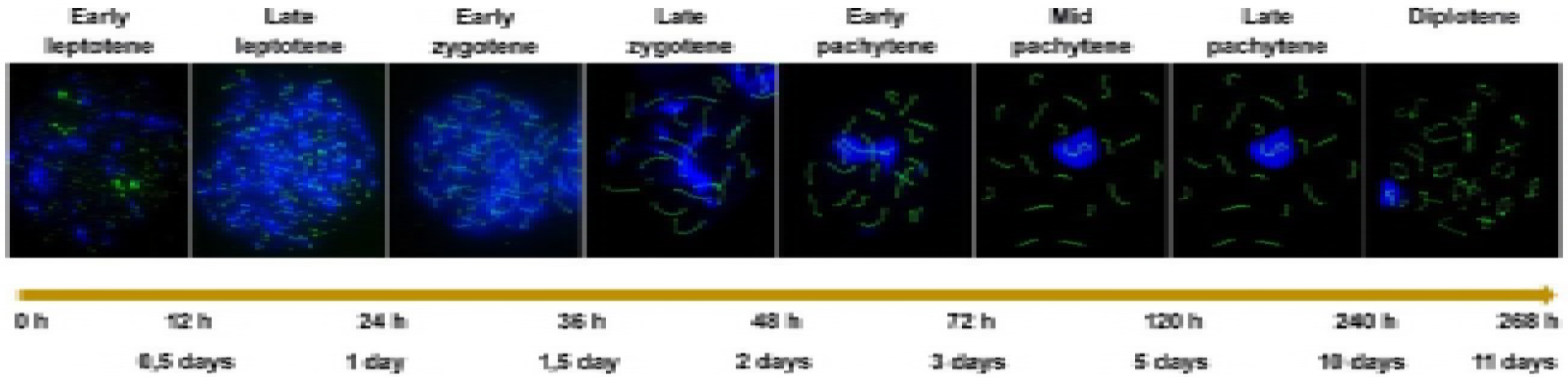
**Estimated length of meiotic stages**, based on previous reports by Oakberg [48] and Ashley and coworkers [47].

**S2 Figure.**
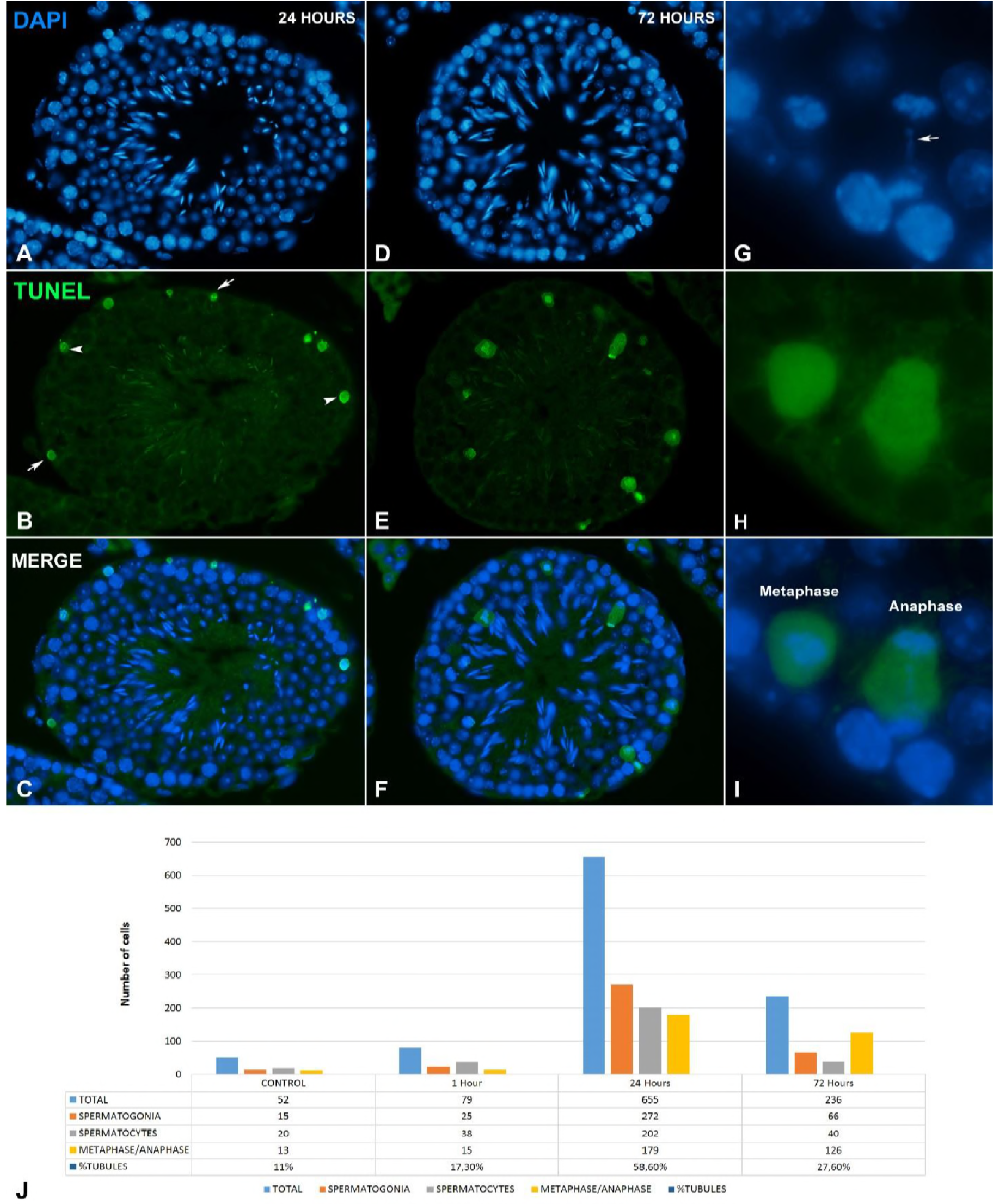
Apoptosis induction after irradiation. TUNEL (green) and DAPI (blue). (A-C) Section of a seminiferous tubule 24 hours after treatment. Apoptotic cells are found in both the basal (arrows) and interstitial (arrowheads) strata of the seminiferous epithelium. (D-F) Section of a seminiferous tubule 72 hours after treatment showing apoptotic cells at metaphase or anaphase. (G-I) Detail of apoptotic metaphase and anaphase cells 72 hours after irradiation. Note the presence of chromatin bridges between cell poles in the anaphase cell (arrow in G). (J). Quantitative distribution of apoptotic cells. Total number of apoptotic cells were recorded in 300 seminiferous tubules. Peak apoptosis is observed 24 after irradiation with 58.6% of tubules showing at least one apoptotic cell. At this time, spermatogonia are the most affected population, followed by spermatocytes and cells undergoing division. After 72 hours of recovery, the total number of apoptotic cells decreases with only 27.6% of tubules showing apoptotic cells. At this recovery time, the majority of cells undergoing apoptosis are at metaphase or anaphase.

**S3 Table.** Quantitative data for all proteins analyzed and chromosomal bridges organized by cell stage and recovery time.

| γH2AX | Mid pachytene |  |  | Late Pachytene |  |  | Early Diplotene |  |  |
| --- | --- | --- | --- | --- | --- | --- | --- | --- | --- |
|  | Mean | SD | n | Mean | SD | n | Mean | SD | n |
| CONTROL | 0.23 | 0.43 | 25 | 0.31 | 0.47 | 25 | 0.08 | 0.27 | 25 |
| 1 HOUR | 50.66 | 4.72 | 29 | 51.46 | 5.89 | 25 | 52.96 | 4.61 | 29 |
| 24 HOURS | 1 | 1.17 | 26 | 7.15 | 2.38 | 26 | 13.31 | 3.89 | 26 |
| 72 HOURS | 0.54 | 0.71 | 26 | 7.35 | 3.59 | 26 | 10.38 | 2.33 | 26 |

| BRIDGES | Early pachytene |  | Mid pachytene |  | Late Pachytene |  | Early Diplotene |  | TOTAL |  |
| --- | --- | --- | --- | --- | --- | --- | --- | --- | --- | --- |
|  | % | n | % | n | % | n | % | n | % | n |
| CONTROL | 0 | 48 | 0 | 101 | 0 | 110 | 0 | 57 | 0 | 316 |
| 1 HOUR | 13.64 | 44 | 5.98 | 117 | 5.93 | 118 | 2.22 | 45 | 6.48 | 324 |
| 24 HOURS | 40.00 | 35 | 6.67 | 120 | 5.98 | 117 | 1.79 | 56 | 9.15 | 328 |
| 72 HOURS | 33.33 | 21 | 30.89 | 123 | 19.40 | 134 | 13.51 | 37 | 24.13 | 315 |

| DMC1 | Early Leptotene |  |  | Mid-late Leptotene |  |  | Early-mi Zygotene |  |  | Late Zygotene |  |  | Early Pachytene |  |  | Mid Pachytene |  |  |
| --- | --- | --- | --- | --- | --- | --- | --- | --- | --- | --- | --- | --- | --- | --- | --- | --- | --- | --- |
|  | Mean | SD | n | Mean | SD | n | Mean | SD | n | Mean | SD | n | Mean | SD | n | Mean | SD | n |
| CONTROL | 7.04 | 5.39 | 24 | 63.62 | 34.49 | 21 | 149.75 | 36.41 | 36 | 111.10 | 19.74 | 29 | 78.89 | 11.08 | 27 | 55.00 | 21.21 | 29 |
| 1 HOUR | 76.08 | 32.15 | 24 | 208.25 | 53.11 | 20 | 226.57 | 32.06 | 30 | 157.50 | 29.62 | 20 | 96.79 | 33.14 | 24 | 58.87 | 19.49 | 31 |
| 24 HOUR | 19.48 | 6.45 | 27 | 162.68 | 47.62 | 28 | 191.70 | 40.12 | 27 | 143.36 | 36.43 | 25 | 100.50 | 29.76 | 26 | 54.96 | 26.56 | 25 |
| 72 HOUR |  |  |  |  |  |  | 188.60 | 42.45 | 25 | 132.62 | 33.07 | 29 | 108.96 | 18.01 | 27 | 55.88 | 22.51 | 25 |

| RAD51 | Mid pachytene |  |  | Late Pachytene |  |  | Early Diplotene |  |  | Late Diplotene |  |  |
| --- | --- | --- | --- | --- | --- | --- | --- | --- | --- | --- | --- | --- |
|  | Mean | SD | n | Mean | SD | n | Mean | SD | n | Mean | SD | n |
| CONTROL | 50.61 | 20.76 | 23 | 0.85 | 1.66 | 25 | 0.23 | 0.65 | 25 | 0 | 0 | 26 |
| 1 HOUR | 56.80 | 14.42 | 27 | 16.62 | 18.06 | 28 | 11.04 | 6.60 | 24 | 6.64 | 3.82 | 30 |
| 24 HOURS | 79.36 | 14.07 | 26 | 87.36 | 18.51 | 23 | 41.08 | 12.18 | 25 | 27.04 | 4.53 | 25 |
| 72 HOURS | 79.88 | 17.95 | 25 | 60.10 | 16.92 | 25 | 18.64 | 5.44 | 26 | 18.42 | 6.00 | 26 |

| RAD51 | Early diplotene ON |  |  | Early diplotene OFF |  |  | Late Diplotene ON |  |  | Late Diplotene OFF |  |  |
| --- | --- | --- | --- | --- | --- | --- | --- | --- | --- | --- | --- | --- |
|  | Mean | SD | n | Mean | SD | n | Mean | SD | n | Mean | SD | n |
| CONTROL | 0.23 | 0.65 | 25 | 0 | 0 | 25 | 0 | 0 | 26 | 0 | 0 | 26 |
| 1 HOUR | 10.00 | 5.78 | 24 | 1.04 | 1.51 | 24 | 5.83 | 3.64 | 24 | 1.08 | 1.41 | 24 |
| 24 HOURS | 36.12 | 8.96 | 25 | 4.96 | 2.46 | 25 | 18.12 | 6.05 | 26 | 8.92 | 4.53 | 26 |
| 72 HOURS | 15.96 | 4.83 | 26 | 2.68 | 2.17 | 26 | 13.42 | 5.30 | 26 | 5.00 | 2.70 | 26 |

| 53BP1 | Mid pachytene |  |  | Late Pachytene |  |  | Early Diplotene |  |  |
| --- | --- | --- | --- | --- | --- | --- | --- | --- | --- |
|  | Mean | SD | n | Mean | SD | n | Mean | SD | n |
| CONTROL | 0.00 | 0.00 | 25 | 0.16 | 0.47 | 25 | 0.50 | 0.91 | 26 |
| 1 HOUR | 27.48 | 5.62 | 27 | 32.85 | 5.89 | 27 | 34.33 | 4.66 | 30 |
| 24 HOURS | 0.48 | 0.71 | 25 | 1.63 | 1.55 | 27 | 4.63 | 2.22 | 27 |
| 72 HOURS | 0.08 | 0.28 | 25 | 0.84 | 1.40 | 26 | 0.81 | 1.27 | 25 |

**S4 Figure.**
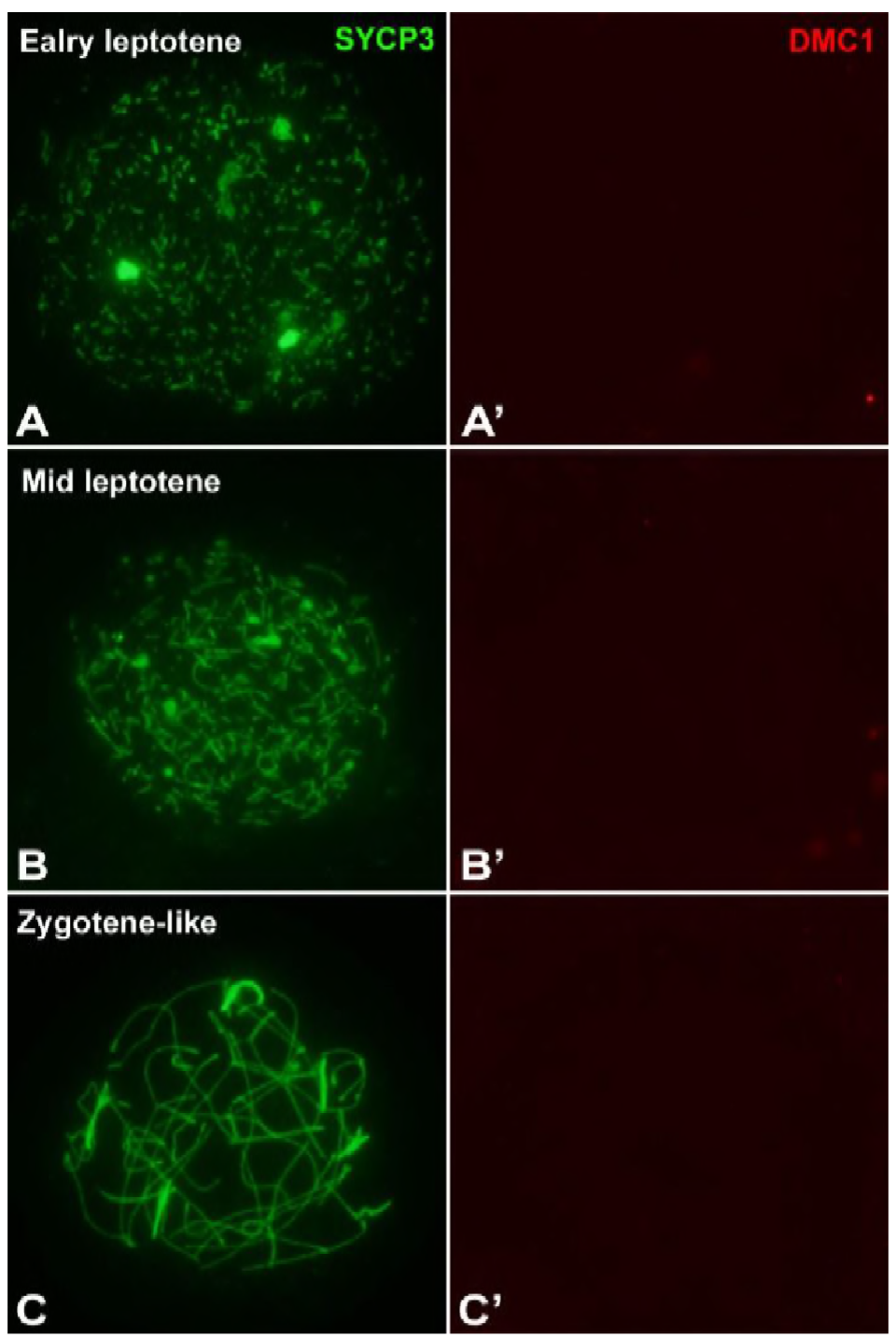
DMC1 localization in SPO11 knockout mice. (A-C) SYCP3 (green) and (A’-C’) DMC1 (red) at early leptotene (A, A’), mid leptotene (B, B’) and zygotene-like (C, C’). No specific signal of DMC1 is detected at any of the stages analyzed.

**S5 Figure.**
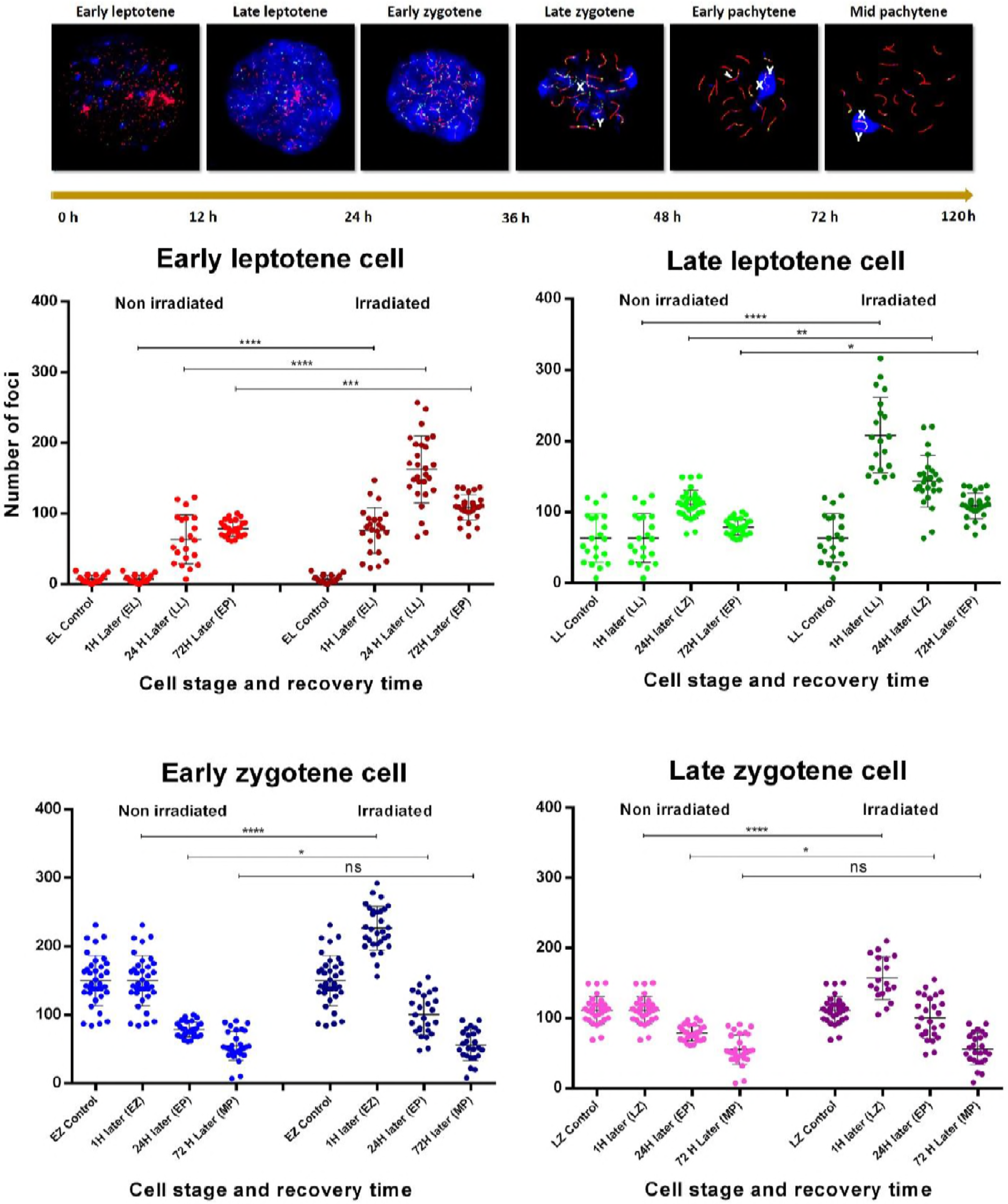
Quantitative analysis of DMC1 dynamics during early prophase-I. Cell progression during prophase-I and duration of each stage is represented in the series of images at the top. Distribution of DMC1 foci are arranged according to the recovery time after irradiation and the putative stages that cells should have reached at that time, provided that meiotic progression was not affected by the treatment. We arranged four cell populations: early leptotene, mid-late leptotene, early-mid zygotene and late zygotene. For each case, irradiated cells were compared with their respective control counterparts and statistical differences indicated (ANOVA and Tukey’s multiple comparisons test). Only cells that advanced to mid pachytene during the recovery time, i.e., those that were irradiated at early or late zygotene, show control levels of DMC1 after 72 hours of recovery. This is mostly due to the fact that DMC1 is not inducible at this stage and/or the programmed displacement of DMC1 from the chromosomes, regardless of whether repair had been completed or not. EL: early leptotene; LL: mid-late leptotene; EZ: early-mid zygotene; LZ: late zygotene; EP: early pachytene; MP: mid pachytene; ns: non-significant; *: p≤0.05; **: p≤0.01; ***: p≤0.001; ****: p≤0.0001.

**S6 Figure.**
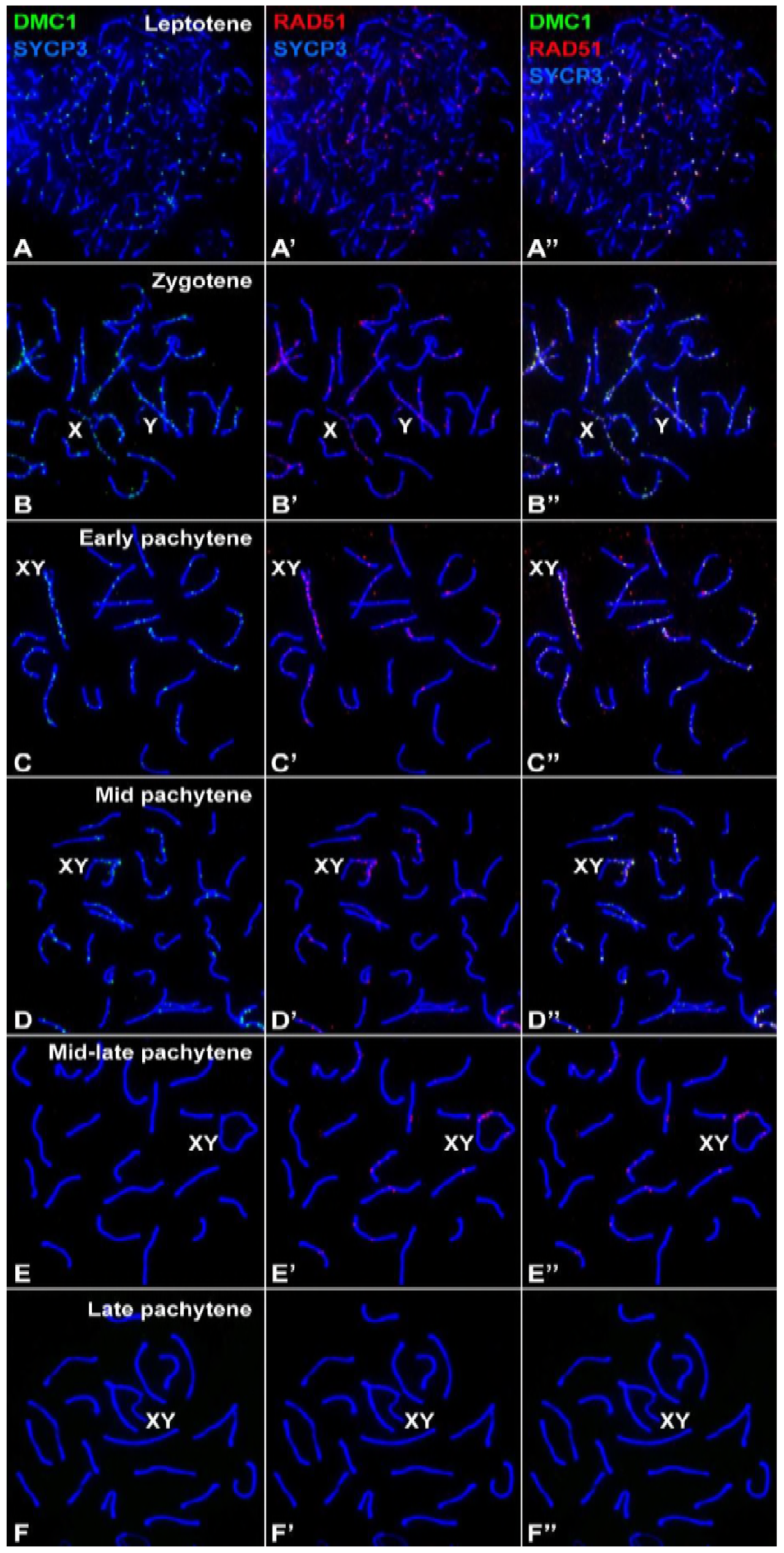
Localization of DMC1 and RAD51 in control spermatocytes. SYCP3 (blue), DMC1 (green) and RAD51 (red). Merge of the (A-F) SYCP3 and DMC1 channels, (A’-E’) SYCP3 and RAD51 channels and (A”-F”) SYCP3, DMC1 and RAD51 channels. DMC1 and RAD51 are largely co-localized in foci observed from early leptotene to mid pachytene (A-D). However, DMC1 and RAD51 signal on these foci are usually not identical in size or shape. Moreover, there are some instances in which either of the two proteins seem to form single foci. At mid-late pachytene (E-E”), DMC1 is no longer present on the chromosomes, but RAD51 is still abundantly observed on both autosomes and sex chromosomes (X and Y). By late pachytene (F-F”), neither DMC1 nor RAD51 are observed.

**S7 Figure.**
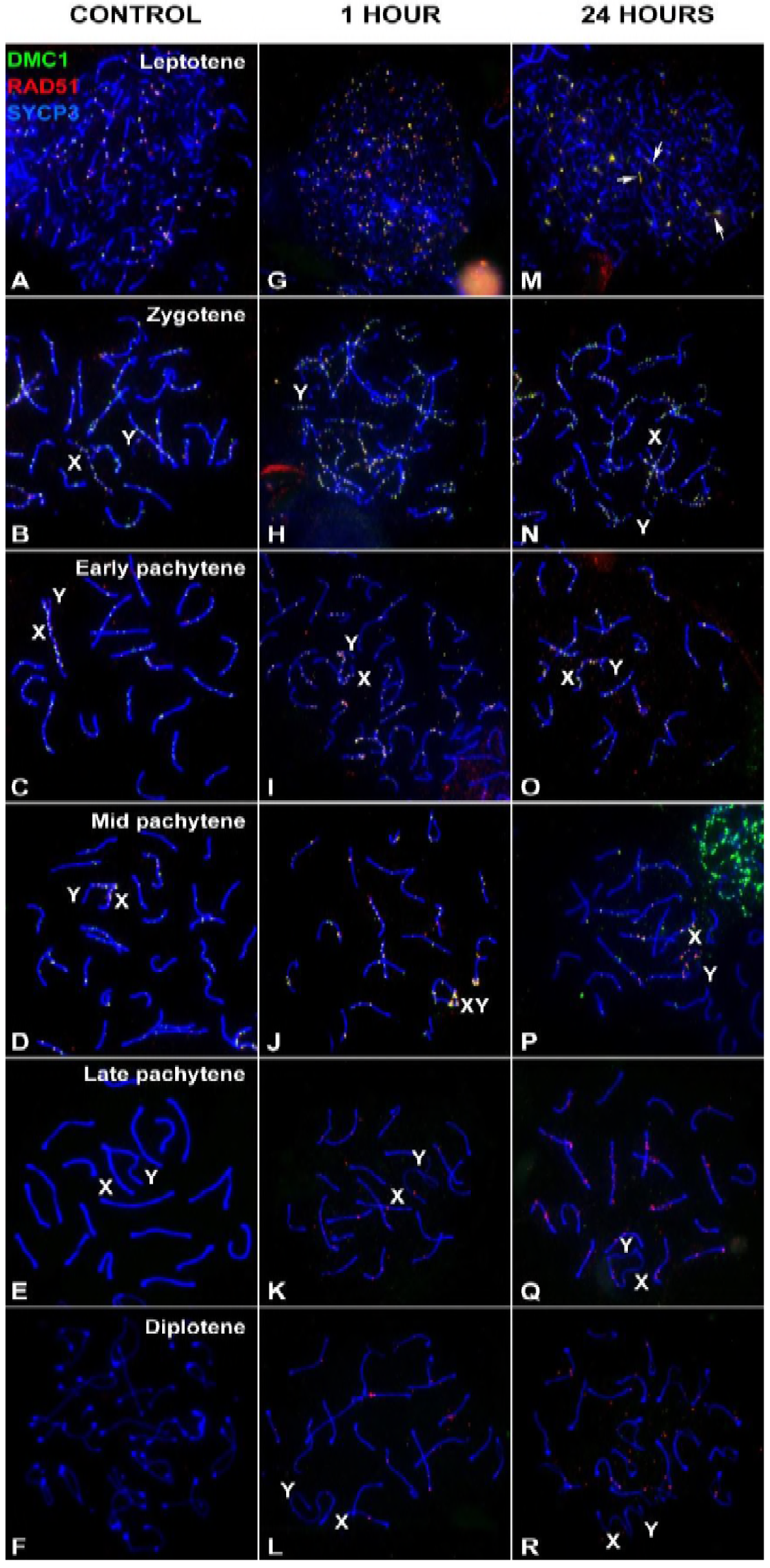
Localization of DMC1 and RAD51 at different stages of prophase-I by different recovery time after irradiation. SYCP3 (blue), DMC1 (green) and RAD51 (red). (A-F) Control. The cells shown in A-D are the same as those shown in S6 Figure. (G-L) 1 hour after irradiation. As shown in Figures 4 and 6, both DMC1 and RAD51 become more abundant after irradiation with both proteins being present in the same foci in most cases; however, the overlap in signals is not identical in many instances as the sizes and shapes of foci of the individual proteins differ. From late pachytene onwards, only RAD51 foci are detectable. (MR) An analogous result is found 24 after irradiation. Filaments containing both DMC1 and RAD are observed in some spermatocytes (arrows in M).

**S8 Figure.**
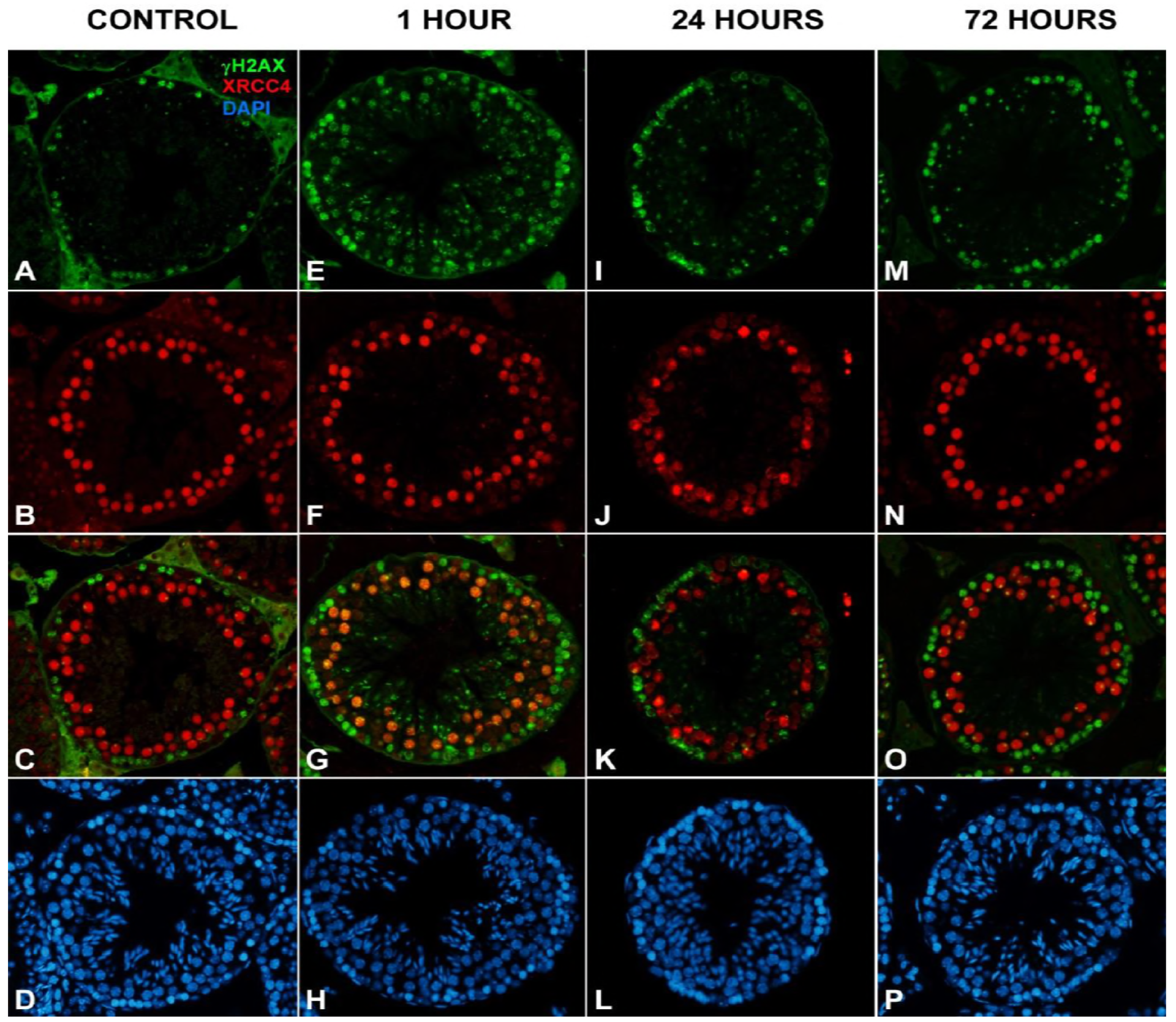
Localization of XRCC4 in testicular sections. γH2AX (green), XRCC4 (red) and DAPI (blue) in seminiferous tubules at equivalent developmental stages. (A-D) Control. Basal layers of spermatocytes, corresponding to leptotene and zygotene, are broadly stained with γH2AX and devoid of XRCC4. Spermatocytes in the interstitial strata of the epithelium show an inverse labeling pattern, with abundant XRCC4 and nearly no γH2AX staining. (E-H). 1 hour after irradiation. Cells showing broad γH2AX labeling in the basal strata are again devoid of XRCC4. In contrast, spermatocytes stained with XRCC4 now also have an abundance of γH2AX localized foci, corresponding to the late γH2AX response. No noticeable increase in the intensity of XRCC4 labeling is observed. (I-L) 24 hours and (M-P) 72 hours after irradiation. γH2AX tend to return to control levels. No variation of XRCC4 labeling is observed after longer periods of recovery.

